# Genome-wide chromatin recording resolves dynamic cell state changes

**DOI:** 10.1101/2025.08.05.668773

**Authors:** Yodai Takei, Jordan A. Lay, James M. Linton, Duncan M. Chadly, Yoshiki Ochiai, Ron Hadas, Andrew A. Perez, Mario R. Blanco, Paola Laurino, Mitchell Guttman, Michael B. Elowitz

**Author notes:** Corresponding author: M.B.E.

## Abstract

Understanding how the chromatin state of a cell influences its future behavior is a major challenge throughout biology. However, most chromatin profiling methods are limited to endpoint assays. Here, we present LagTag, a method for recovery of earlier and endpoint chromatin states in the same mammalian cells. In this approach, transient expression of bacterial adenine methyltransferase fusions records the DNA binding profiles of chromatin-associated proteins of interest at earlier timepoints. Subsequent tagmentation and sequencing recovers the earlier chromatin profile from adenine methylation profiles, alongside endpoint profiles of endogenous chromatin-associated proteins. We verified that LagTag profiles aligned with those from established methods in mouse and human cells. We then applied LagTag to record and recover dynamic chromatin state transitions during mouse embryonic stem cell differentiation, capturing transcriptional signatures from pre- and post-differentiation timepoints within the same cell population. LagTag thus provides a foundation for temporally resolved chromatin profiling.

## Introduction

Chromatin states powerfully influence subsequent gene regulation and cellular behavior. They impact signal responses, cell fate decisions, and disease progression^1^. Knowing the causal relationships between earlier chromatin states and subsequent cellular behaviors is therefore critical for understanding, predicting, and controlling gene regulation and cell states. Existing approaches for mapping chromatin states generally provide static “snapshots” from single endpoint measurements^2^, providing no direct readout of earlier states. The ability to recover information from multiple timepoints in the same cells would reveal dynamic relationships that are otherwise difficult to infer. Temporally integrated measurements could reveal if and when pre-existing chromatin states influence cell fate decision-making. They could also identify chromatin states that lead to cancer drug tolerance^3^. An additional advantage of chromatin-based recording is its ability to capture various repressive chromatin states^4^ at transcriptionally inactive loci, information that is not attainable through transcription-based recording alone^5,6^.

Recent work has attempted to address this fundamental challenge in different ways. Pseudotime approaches can infer likely developmental trajectories of individual cells based on cell state similarity in static snapshots^7^. However, these approaches do not provide information from multiple timepoints in the same cell^8^. RNA velocity can provide insights into the instantaneous rates of change of RNA species in a cell^7,9^, but this is distinct from capturing a specific earlier timepoint and can be unreliable^10^. In the Rewind method^11^, inherited barcodes allow one to compare cells at a later timepoint to the state of their siblings at an earlier timepoint. However, this approach is challenging to apply in tissues as it requires splitting cell populations, and assumes strong sister cell similarity, which may not always apply. A parallel set of synthetic biology approaches record information by continuously editing genomically integrated barcodes in a manner that allows reconstruction of lineage and signaling events^12^. However, these approaches rely on cis-regulatory elements engineered to respond to signals of interest and do not directly provide information about chromatin states across the genome. Finally, other approaches such as engineered RNA exporters^6^ or Live-seq^5^ have been developed to allow non-destructive measurements of RNA over time in the same cells, but do not directly provide information about chromatin states. Thus, the ability to recover chromatin state dynamics across the genome at multiple timepoints has remained a challenge.

Some chromatin profiling approaches store chromatin state information in live cells, potentially enabling dynamic measurements. In particular, while adenine DNA methylation rarely occurs in mammalian cells naturally, it can be implemented through recruitment of bacterial adenine methyltransferases to DNA in live cells^13^. More specifically, by fusing a protein of interest with bacterial DNA adenine methyltransferase (Dam), the DamID method successfully recorded genome-wide DNA-protein interactions through profiles of adenine DNA methylation at GATC motifs^14^. MadID enhanced this approach, using a non-specific adenine methyltransferase (M.EcoGII) that does not require a specific consensus sequence to provide broader genome coverage^15^. The same work also showed that antibodies against methylated adenine (m6A) could be used in place of methylation-sensitive restriction enzymes (e.g. DpnI) to recognize adenine methylation (m6A) independent of sequence motifs. Finally, in the absence of active adenine methylation, deposited m6A modifications are partitioned to daughter cells^16^. Thus, m6A marks can be “written” in living cells, read out in fixed cells, and diluted, but not erased, during cell division. However, despite the potential of chromatin recording in live cells, these genome-wide approaches have primarily been applied to recording endpoint chromatin states by expressing fusion proteins just before harvesting cells. It has remained unclear whether they can also enable recording of past genome-wide chromatin states.

Recently, two chromatin recording approaches^17,18^ were introduced, both of which take advantage of bacterial methyltransferases to record information in DNA modifications. DCM-time machine (DCM-TM)^17^ uses the bacterial methyltransferase DCM fused to an RNA polymerase II subunit to label active genes and enhancers with cytosine methylation at specific sequence motifs. This approach revealed dynamic chromatin state changes during enterocyte differentiation in the mouse intestine. DCM methylation profiles were recorded in different time windows in different samples. However, this approach revealed only recorded chromatin profiles and could not also reveal endpoint chromatin profiles from the same cells. In contrast, the Dam&ChIC approach^18^ enabled temporal chromatin profiling in single cells by measuring past chromatin profiles recorded by Dam together with present chromatin profiles by ChIC simultaneously. This approach resolved temporally ordered chromatin events such as nuclear lamina detachment and H3K27me3 accumulation during X chromosome inactivation^18^. However, the DpnI restriction enzyme used in DamID cleaves fully methylated DNA much faster than hemi-methylated DNA containing the GATC motif^19^. As a result, its operation is typically limited to a time interval less than one cell cycle^16,18^. Thus, despite much work, a method that allows chromatin recording of states further into the past alongside endpoint chromatin measurements has been lacking.

Here we present LagTag, a technology that records chromatin states by pulse-labeling with m6A and recovers them alongside additional chromatin features by simultaneous readout at a later endpoint. m6A recording performed immediately before the endpoint captures the present chromatin states, similarly to conventional DamID approaches, whereas a subsequent chase period after recording enables retrospective profiling of past chromatin states across cell divisions. Technically, LagTag combines the temporal recording capabilities of bacterial methyltransferases in live cells with post-fixation tagmentation and sequencing readout. The use of tagmentation substantially reduces cell input requirements compared to immunoprecipitation-based methyltransferase methods such as MadID^15^. Importantly, LagTag extends the temporal recording window of DamID to capture chromatin states separated by at least two cell cycles. LagTag effectively generates historical records of chromatin states that can be validated against established measurements such as those from ChIP-seq^20^ and CUT&Tag^21^. We first establish the technical foundation of LagTag (**Figure 1**) and then present proof-of-principle experiments demonstrating that temporal chromatin recording captures earlier chromatin signatures during mouse embryonic stem cell differentiation to endodermal fates^22^ (**Figure 2**).

**Figure 1.**
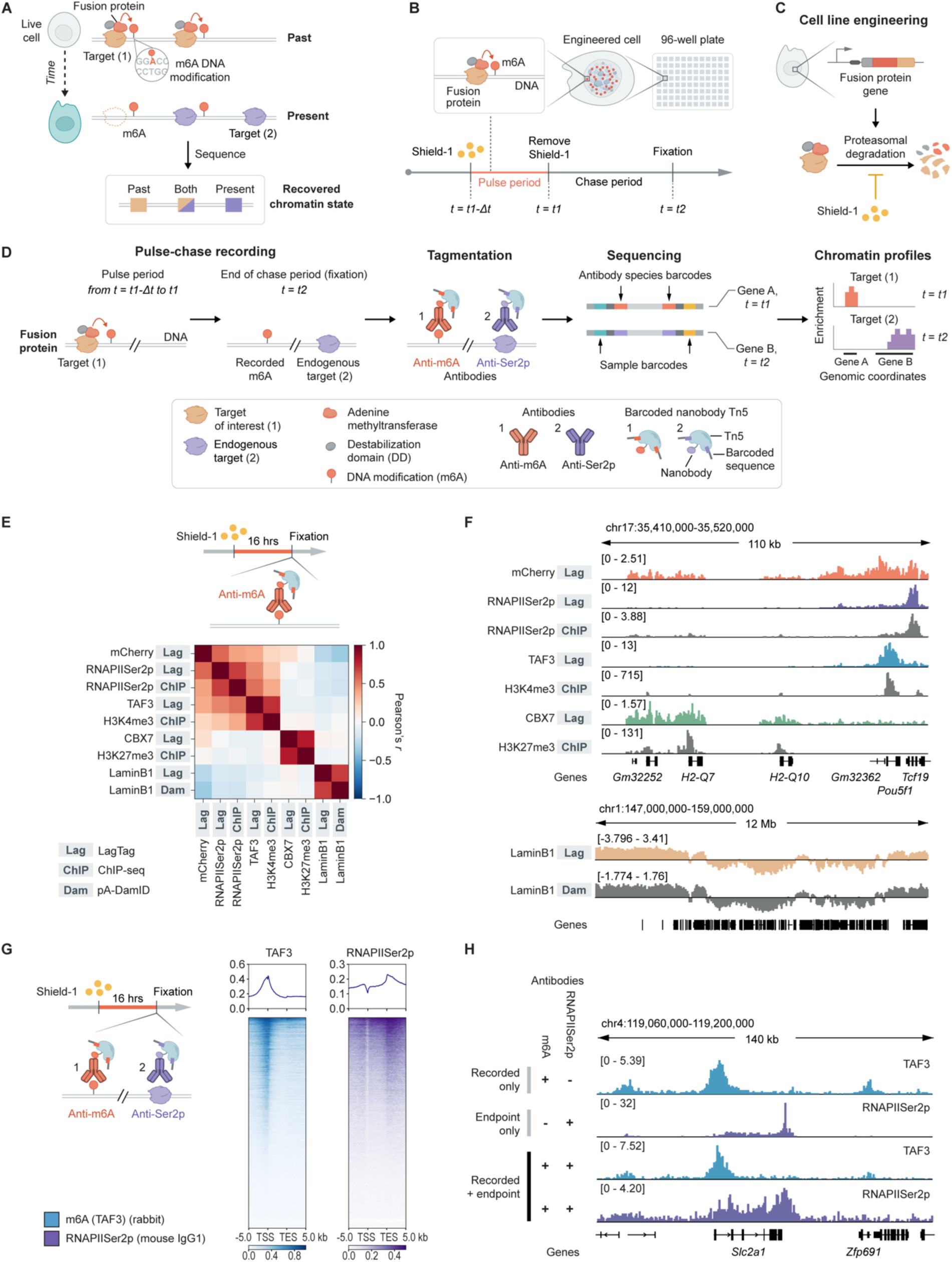
LagTag platform includes a temporal recorder and endpoint readout. **A**, The principle of recovering earlier chromatin states via recorded m6A DNA methylation alongside present states. **B**, **C**, Schematic illustration of the temporal recording of chromatin states. In engineered cell lines, the fusion protein between protein of interest and adenine methyltransferase controls the deposition of m6A DNA methylation based on its protein-DNA interactions during the pulse period (**B**). The small molecule Shield-1 can modulate the stability of the fusion protein (**C**) to achieve temporal chromatin recording. **D**, Schematic representation of the LagTag procedure, including chromatin state recording in live cells, recovery of recorded states via antibody-guided tagmentation in fixed cells, sequencing, and downstream analysis. Target of interest (1) during the recording and endogenous target (2) at the endpoint (e.g., RNAPIISer2p) can correspond to either the same or different molecular species. **E**, Schematic illustration of LagTag single-antibody assays (top). Correlation heatmaps of chromatin profiles for different targets as well as different profiling methods in mESCs (bottom). Correlations were calculated across 10-kb bins that overlapped with the union of peaks previously called from the ChIP-seq (H3K4me3 and H3K27me3) and CUT&Tag (RNAPIISer2p) datasets (n = 61,536 bins). **F**, Genome browser tracks of LagTag, ChIP-seq^20,37^, and pA-DamID^33^ samples in mESCs. Scale of LagTag profile was normalized to coverage per million aligned reads (top) or normalized to log2 ratio between Lamin B1 and mCherry profiles (bottom). The ChIP-seq profiles are input normalized and pA-DamID profiles are normalized by the Dam only control. n = 2 for TAF3 and CBX7, and n = 3 for mCherry, RNAPIISer2p, and Lamin B1 in LagTag experiments in **E** and **F**. **G**, Schematic illustration of LagTag multi-antibody assays (left). Signal enrichment plots for demultiplexed LagTag profiles for two antibodies around transcription start site (TSS) and transcription end site (TES) across all protein-coding genes in mESCs (right). **H**, Genome browser tracks of demultiplexed LagTag profiles for two antibodies along with individual LagTag and CUT&Tag profiles with single antibodies in mESCs. Normalization of the scale was performed similarly to **F** for LagTag and CUT&Tag. n = 2 for experiments in **G** and **H**.

**Figure 2.**
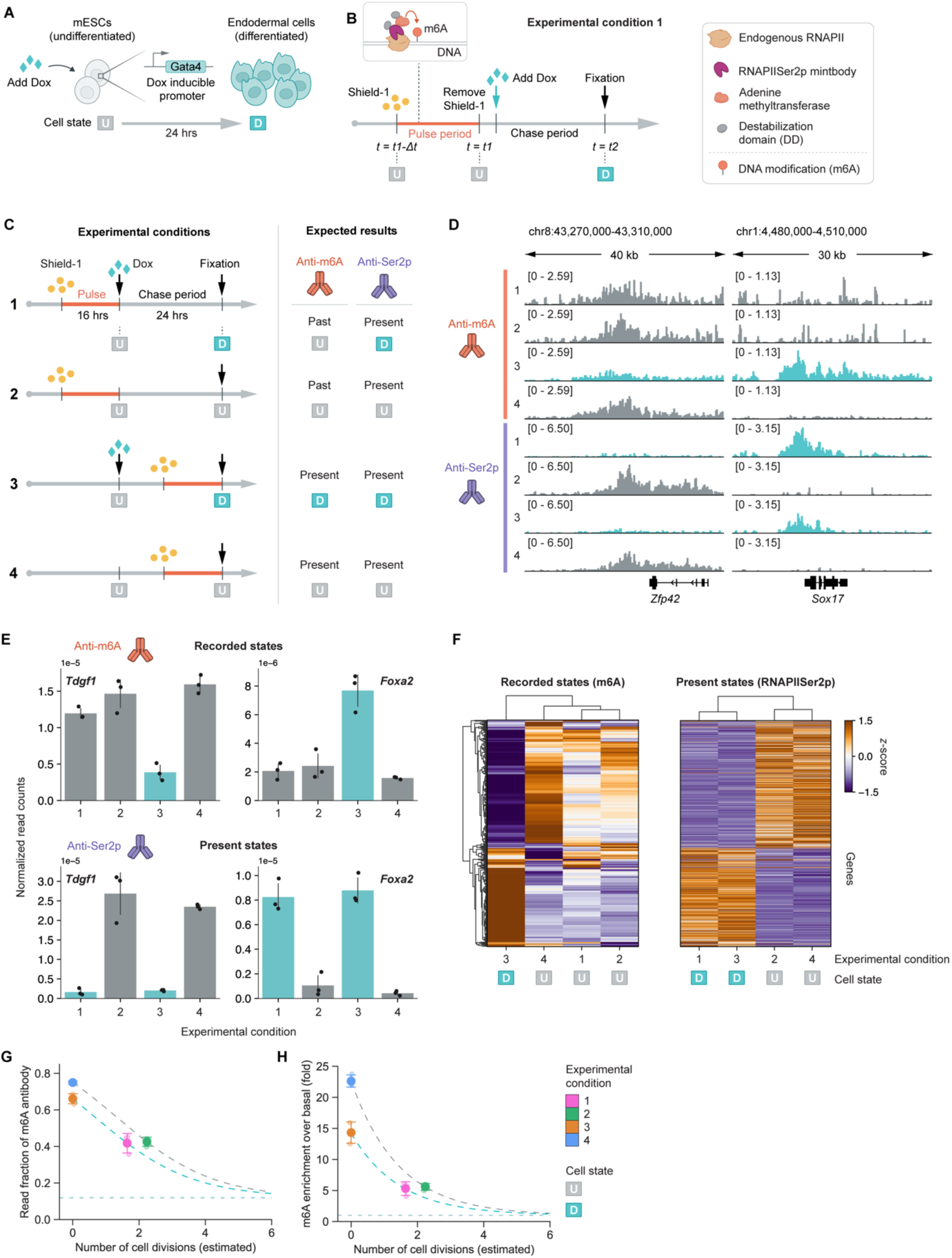
LagTag enables simultaneous chromatin profiling of two timepoints from the same samples. **A**, Schematic illustration of cellular differentiation in mESCs with inducible Gata4 system. **B**, LagTag allows temporal chromatin profiling by recording the previous chromatin states (undifferentiated cell state, U) and reading out after mESC differentiation (differentiated cell state, D). **C**, Additional schematic representation of LagTag experimental conditions with mESC differentiation along with expected results, corresponding to the recorded and endpoint chromatin states in each condition. The more detailed schematic for experimental condition 1 is depicted in **B**. **D**, Genome browser tracks of recorded and endpoint LagTag profiles across experimental conditions. **E**, Comparison of normalized read counts across experimental conditions. The color bars are colored by the expected cell states. The error bars represent standard deviations. **F**, Cluster maps of genes and conditions for recorded (left) and present (right) chromatin states using the differentially enriched genes with RNAPIISer2p (n = 933 genes) between undifferentiated and differentiated cell states, defined by separate CUT&Tag experiments (n = 3). Gene ordering in the right panel follows the clustering in the left panel. **G**, **H**, Read fraction of the m6A antibody (**G**) or m6A fold enrichment over basal (**H**) as a function of estimated cell divisions, modeled using a cell division dilution framework (see **Methods**). Error bars represent mean ± standard deviations with individual replicate points overlaid. Results are shown separately for undifferentiated and differentiated cell states. Basal levels (dashed asymptotes) were determined from uninduced controls (conditions 5 and 6). Dashed curves show the predicted decay over cell divisions, derived from the measured basal and induced conditions (see **Figure S9B**). n = 3 for LagTag experiments in **D**-**H**.

## Results

### LagTag combines m6A recording with tagmentation-based endpoint readout

To develop m6A-based chromatin recording, we combined MadID^15^ and CUT&Tag^21^, which provide complementary capabilities. MadID deposits m6A modifications at specific genomic regions bound by the methyltransferase fusion to enable recording^15^, but uses immunoprecipitation, which typically requires millions of cells, for sequencing readout. By contrast, CUT&Tag does not record earlier events, but enables chromatin profiling of multiple chromatin marks and proteins in fewer, or even single, cells at the endpoint by recruiting transposase to an antibody of interest bound at specific chromatin sites^21,23–26^. We reasoned that these approaches could be combined using m6A deposition to record chromatin states and antibody-guided tagmentation to read them out (**Figure 1A**). We term this combined approach LagTag, as it enables later (time-lagged) recovery of earlier chromatin profiles.

More specifically, for recording, we constructed fusions of the M.EcoGII adenine methyltransferase to chromatin binding proteins, including full-length proteins, their domains, and modification-specific intracellular antibodies (mintbodies)^27,28^ (**Figure S1A**), and minimized expression levels via a mutated Kozak sequence with reduced translation efficiency^29^. To sharply define the temporal recording window, we also incorporated a destabilization domain (DD) that can be inhibited by the Shield-1 small-molecule inducer^15,16,30^. With this modular system, adding and subsequently washing out Shield-1 leads to transient genome-wide adenine methylation at the specific sites where the protein of interest is localized during the labeling phase (**Figure 1B-D**). Recorded m6A marks can then be read out by sequencing via antibody-guided tagmentation using an m6A antibody^15^. Endpoint chromatin profiles can be read out using other CUT&Tag compatible antibodies. In particular, using a second, orthogonal antibody against elongating RNA polymerase II marked by Serine 2 phosphorylation (RNAPIISer2p) provides a readout of active expression at the final timepoint (**Figure 1D**). We note that the engineered fusion protein used to record the past state can be the same as or different from the endogenous target detected in the present state, as they are independent.

### LagTag reproduces known chromatin profiles

To validate LagTag, we first asked whether recorded m6A profiles aligned with publicly available datasets in mouse embryonic stem cells (mESCs) and human HCT116 cells. To record, we fused the non-specific adenine methyltransferase M.EcoGII^15^ with different full-length proteins, protein domains, and mintbodies (**Figure 1B, Figure S1A**), then generated stable cell lines expressing each construct individually (**Figure 1C**). First, to target endogenous elongating RNAPII, we fused M.EcoGII and DD with a mintbody targeting RNAPIISer2p (Ser2p-mintbody)^31^. Second, to profile histone modifications, we fused M.EcoGII and DD with well-characterized chromatin reader domains from TAF3 and CBX7, which recognize H3K4me3 and H3K27me3, respectively^27,28^. Third, to mark repressive lamina-associated domains (LADs), we fused M.EcoGII and DD to full-length Lamin B1, a protein that binds to the nuclear lamina and is widely used for DamID and related approaches^15,16,32^. Finally, as a negative control, we also fused M.EcoGII and DD to a non-specific mCherry fluorescent protein. Together, these constructs provided tools for inducible temporal control of m6A recording of chromatin states.

Using these constructs, we induced recording for 16 hours in mESCs and HCT116 cells grown in standard media in individual wells of 96-well plates containing ∼10,000-30,000 cells (see **Methods**). We then fixed cells with 1% formaldehyde for 5 minutes, performed endpoint assays, and compared the results with existing data sets (**Figure 1E, F, Figure S1-4, Table S1, 2**). In mESCs, the Ser2p-mintbody, TAF3 and CBX7 reader domains, Lamin B1, and mCherry (negative control) all exhibited distinct genomic enrichment patterns, as expected (**Figure 1E, F, Figure S2**). Specifically, mCherry, Ser2p-mintbody, and TAF3 were associated with active chromatin, CBX7 was associated with facultative heterochromatin, and Lamin B1 was associated with repressive LADs. Further, these profiles agreed with independent chromatin profiling measurements using ChIP-seq^20^, CUT&Tag^21^, and pA-DamID^33^. For example, in mESCs, the recorded TAF3 (H3K4me3 reader domain) profiles were similar to those obtained by ChIP-seq and CUT&Tag (Pearson’s correlation coefficients of 0.76 and 0.87, respectively for H3K4me3, **Figure 1F, Figure S2C, E, F**). The LagTag profiles also showed specific enrichment patterns at transcription start sites (TSSs) for TAF3, transcription end sites (TESs) for RNAPIISer2p^34^, or called H3K27me3 peaks for CBX7 (**Figure S3A**). Similarly, background-normalized LagTag and pA-DamID signals were strongly correlated for Lamin B1^33^ (**Figure 1F, Figure S2D, F**, Pearson correlation coefficient of 0.91). We note that the modified CUT&Tag protocol used here efficiently recovered tagmented fragments from fixed cells (**Figure S1B**) and enabled mapping of Lamin B1 (**Figures S2D-F, 3B**), which, until recently, was considered difficult to profile by tagmentation^35^. Furthermore, we observed good agreement between m6A recorded profiles and independent measurements for RNAPIISer2p and Lamin B1 in human HCT116 cells (**Figure S4**). These data confirm that LagTag, by using m6A marks and an m6A antibody, replicates established measurements that use restriction enzymes or target-specific antibodies (**Figure 1E, F, Figures S2-4**).

One potential limitation of the use of M.EcoGII is its ability to methylate RNA substrates^36^. To identify alternative enzymes that would potentially exhibit reduced RNA methylation, we scanned a large number of M.EcoGII variants (**Figure S5A, B**). LagTag also worked with one of these variants, EspM, a non-specific adenine methyltransferase (**Figure S5C, D**). EspM exhibits a reduced methylation rate for RNA substrates in vitro, while preserving DNA methylation (**Figure S5B**). While M.EcoGII fusion proteins were used for the rest of this study, this result suggests that LagTag can be extended to other adenine methyltransferase variants with distinct properties. Taken together, these results demonstrate that LagTag can accurately profile various active and repressive chromatin regions in mouse and human cell lines at relatively low (∼10,000) numbers of cells.

### LagTag enables simultaneous profiling of m6A-recorded and endpoint chromatin signatures

To obtain chromatin profiles of the same cells at different timepoints, it is necessary to simultaneously read out signatures corresponding to both past and present states (**Figure 1A, D**). To test this ability, we adapted multifactorial CUT&Tag approaches previously used to map multiple chromatin proteins and modifications in single assays^23–26^. Specifically, we adapted nanobody-based multifactorial CUT&Tag, which employs uniquely barcoded nanobody-transposases (nb-Tn5) to orthogonally bind to and distinguish the different antibody species (i.e. rabbit and mouse IgG1)^25,26^ (**Figure S6A**).

Using the optimized 96-well format described above, multifactorial CUT&Tag worked robustly across multiple targets, including RNAPIISer2p, H3K4me3, and H3K27me3 in fixed cells (**Figure S6A, B**). For example, consistent with previous studies, the profiles of RNAPIISer2p and H3K27me3 showed minimal overlap in mESCs^23^ (**Figure S6C-E**). This indicates there is minimal cross talk between barcodes targeting different molecular species.

Next, we performed LagTag assays using two antibodies (e.g., m6A and RNAPIISer2p) and the two barcoded nb-Tn5 species together on the same cell population (**Figure 1G, H, Figure S7A-F**). The combined assays produced similar chromatin profiles as the individual LagTag and CUT&Tag assays for TAF3 and RNAPIISer2p pairs (Pearson correlation coefficients of 0.98 and 0.81) as well as Lamin B1 and RNAPIISer2p pairs (Pearson correlation coefficients of 0.96 and 0.81) (**Figure S7C-E**). We also confirmed the expected differential enrichment of TAF3 and RNAPIISer2p at TSS and TES, respectively, as well as Lamin B1 enrichment and RNAPIISer2p depletion at LADs (**Figure 1G, H, Figure S7F**). LagTag requires denaturation of chromatin by NaOH in order to enable the m6A antibody to access DNA modifications^15^. This treatment preserved CUT&Tag profiles for RNAPIISer2p at the TES and gene body, enabling the identification of genes with high levels of RNAPIISer2p at their gene bodies (**Figure S7G, H**). However, it also depleted the previously observed RNAPIISer2p enrichment at TSS^37^ for unknown reasons (**Figure S7G**). Together, these results demonstrate simultaneous mapping of m6A-recorded and endpoint chromatin profiles, a critical requirement for LagTag.

### LagTag resolves distinct past and present chromatin states during cellular differentiation

Having established the ability of LagTag to recover recorded and endpoint chromatin profiles, we next asked whether we could use LagTag to analyze dynamic chromatin changes. Cellular differentiation provides an ideal use case, since the influence of earlier chromatin states on subsequent cell state changes is of broad interest but difficult to infer from static snapshots. As a model system, we focused on endodermal differentiation, a key step in many regenerative medicine approaches^38^. We used an mESC line with a doxycycline (Dox)-inducible Gata4 gene, whose expression is sufficient to induce endodermal fate^22^. To enable recording, we stably incorporated a Shield-1-inducible M.EcoGII-Ser2p-mintbody in this cell line. The resulting system produced endodermal differentiation in cell populations in response to 24 hours of Dox exposure in both the parental and M.EcoGII-Ser2p-mintbody engineered cell lines (**Figure S8**). Further, in response to Dox, chromatin profiles of endodermal and pluripotent marker genes switched markedly, with minimal signal remaining from the prior undifferentiated state, suggesting homogeneous changes across the population (**Figure S8C**).

To ensure that recording could be limited to desired time windows, we fused destabilization domains (DDs) to both N- and C- termini of M.EcoGII-Ser2p-mintbody (following a strategy used in ref.^39^). The dual DDs suppressed basal activity, as shown by the increased fold change of m6A marks upon addition of Shield-1 compared to the construct only with the N-terminus DD (**Figure S1F-H**).

Using this engineered cell line, we asked whether it is possible to simultaneously read out the pre- and post-differentiation chromatin profiles of the same cell population. We performed pulse-chase m6A recording by inducing M.EcoGII-Ser2p-mintbody in undifferentiated cells for 16 hours to record chromatin state (pulse) (**Figure 2A, B, Figure S9, 10**). After washing out Shield-1 for a few hours to halt recording, we induced Gata4 for 24 hours to stimulate endodermal differentiation (chase). During this time, cells underwent approximately two cell divisions (2.1 ± 0.3 and 1.9 ± 0.2 divisions, mean ± standard deviation, for parental and engineered cells, respectively) (**Figure S10A, B**). This is consistent with cell cycle times of 13.25 hours (undifferentiated) and 10.5 hours (early-differentiated) reported by live-cell imaging^40,41^.

Following the pulse-chase experiment, we examined the chromatin profiles of pluripotency and differentiation-associated genes (**Figure 2B, C**, condition 1, **Figure S9**). Recorded m6A marks were enriched around the TES of Zfp42, a pluripotency factor expressed in undifferentiated mESCs^42^, reflecting the higher expression of this gene in the past state (**Figure 2C, D**, condition 1). In the same sample at the endpoint, RNAPIISer2p was enriched around the TES of Sox17, a marker gene for endodermal differentiation^22^ (**Figure 2C, D**, condition 1). Conversely, we did not observe enrichment of Sox17 in the recorded m6A profile nor enrichment of Zfp42 in the endpoint RNAPIISer2p profile. These recorded and endpoint chromatin profiles reflect the expected changes of RNAPIISer2p occupancy during differentiation based on our earlier analysis (**Figure S8**).

We also quantified signal enrichment over gene bodies. We confirmed RNAPIISer2p enrichment in the undifferentiated marker gene Tdgf1 in the recorded state and the differentiated marker gene Foxa2 in the endpoint state (**Figure 2E**, condition 1, **Figure S9E**), as expected^22,43^. Finally, we analyzed 933 genes that were differentially enriched with RNAPIISer2p after differentiation (**Figures S8B, S9F**). The recorded m6A signatures from experimental condition 1 clustered together with those from other undifferentiated conditions obtained at either early or endpoint timepoints (**Figure 2F**, left). Similarly, the endpoint RNAPIISer2p signatures from condition 1 clustered together with the other differentiated condition (**Figure 2F**, right), as expected. Together, these results demonstrate that LagTag can recover undifferentiated (past) and differentiated (present) chromatin profiles from the same sample, exhibiting expected differences in specific marker genes and genome-wide profiles.

To confirm these results arose from biological processes rather than technical artifacts, we included several control conditions (**Figure 2C, Figure S9A-D**). First, we omitted Dox to eliminate differentiation and maintain cells in the undifferentiated state (**Figure 2C**, condition 2). When we performed the LagTag assay in this condition, we observed similar profiles between the m6A and RNAPIISer2p marks on marker genes (**Figure 2D, E**). Second, we recorded chromatin states at the endpoint by delaying induction of the M.EcoGII fusion until the last 16 hours before fixing the cells, with or without Dox (**Figure 2C**, conditions 3 and 4). Again, the m6A and RNAPIISer2p profiles corresponding to the same cell state displayed similar trends on marker genes (**Figure 2D, E**), indicating that the assays are largely consistent with one another regardless of cell state and timing of recording. We also extended this comparison to the fold-change across all coding genes during differentiation, confirming that LagTag shows a positive correlation between m6A and RNAPIISer2p (Pearson correlation coefficient of 0.71) as well as in comparison with conventional RNAPIISer2p CUT&Tag (0.65 and 0.76 for m6A and RNAPIISer2p, respectively) (**Figure S9G**). In contrast, changes of some of the shorter non-coding genes (e.g., miRNA) were not detected by LagTag (**Figure S9H**), which represents a potential limitation of the protocol. Third, we included additional negative controls with no expression of the M.EcoGII fusion protein, also with or without differentiation (**Figure S9A**, conditions 5 and 6). When the methyltransferase was not induced (no recording), most sequencing reads were from the endpoint assay of RNAPIISer2p, indicating that LagTag has a low background of basal enzymatic activity (**Figure S9B**). Furthermore, we confirmed that LagTag recording minimally perturbs cellular physiology during differentiation, as measured by cell viability, cell cycling, and instantaneous transcriptional profiles mapped by RNAPIISer2p enrichment in comparison with parental cells (**Figure S10**).

### LagTag resolves past chromatin states within two cell divisions

Finally, we estimated the m6A signal decay as a function of cell divisions (**Figure 2G, H**) by leveraging the read count ratio between m6A and RNAPIISer2p across conditions in combination with a cell division dilution model (see **Methods**). Based on this analysis, we estimate approximately two cell divisions elapsed for undifferentiated and differentiated cells (2.2 ± 0.1 and 1.7 ± 0.3 with mean ± standard deviation, respectively) during the 24-hour chase period, consistent with the direct results from the cell division assay (2.0 ± 0.1 and 1.9 ± 0.2 divisions, respectively) (**Figure S10B**). This model further defined an effective window of approximately two or three cell divisions, within which m6A signal enrichment remains robust (e.g., 14.2 and 4.3 fold m6A signal enrichment over basal in differentiated cells with no chase and 2-division chase, respectively), before progressively approaching background levels after approximately six cell divisions (**Figure 2G, H**). These results establish that LagTag can retrospectively profile chromatin states across at least two cell divisions in proliferating cells.

Taken together, these results demonstrate how LagTag enables genome-wide analysis of past and endpoint chromatin states within the same cell population across cell cycles.

## Discussion

Recovering historical events in cells—especially chromatin states—is essential for understanding how cells change and make fate decisions over time. Compared to previous approaches, LagTag introduces several innovations. First, it extends the duration over which recorded information can be recovered. With conventional DamID, a single round of DNA replication can limit signal recovery, due to the faster digestion of fully methylated DNA compared to hemimethylated by DpnI^19^. Here, by contrast, the use of the m6A antibody inherently allows detection of hemi-methylated strands, which overcomes this limitation, and allows m6A to be used for genome-wide chromatin recording. Second, LagTag enables recovery of endpoint states alongside earlier recorded states in the same cell population. This is made possible by combining m6A for recovery of past information with another antibody (e.g. RNAPIISer2p) capturing the endpoint information. Third, LagTag makes it easier to profile diverse conditions by dramatically decreasing the number of cells required for analysis compared to MadID^15^. Where MadID requires millions of input cells, LagTag requires only ∼10^4^ cells and can therefore be performed in individual wells in 96-well plates. Thus, LagTag allows analysis of two timepoints in a high-throughput fashion with relatively low cell input.

LagTag should address previously intractable biological problems. For example, the origins of heterogeneity in cellular responses are often mysterious^44^. Different cells can exhibit dramatically different responses to signals or drugs, or make distinct cell fate choices in the same environment. The ability to recover earlier chromatin states, even from bulk populations, will help identify signatures that predispose cells to different outcomes or future trajectories in healthy and disease contexts. This is conceptually analogous to two-timepoint mRNA profiling, which has recently enabled transcriptional recording at the population level and identified molecular signatures of drug-naïve persister cells in a human cancer line^45^. Another set of questions pertains to dynamic changes in chromatin state in the central nervous system. Previous work has implicated chromatin changes during memory formation and recall in neurons in the hippocampus^46^. In addition, changes in the subnuclear localization of DNA loci are known to functionally affect transcriptional activation and repression^47^. LagTag could be used to record and recover earlier subnuclear localization states to explore the dynamics and functional impact of nuclear compartment switching during development and aging. While LagTag and other two-timepoint approaches alone do not establish direct causality between earlier chromatin states and subsequent cell fate transitions, they can reveal candidate molecular signatures and temporal ordering of events that are typically inaccessible from static, single time-point measurements^8,18^.

LagTag could be extended in several ways. First, because CUT&Tag and related methods have been shown to operate at the single cell level^21,23–26^, it should be possible to similarly scale down LagTag to access single-cell histories. Single-cell LagTag would enable retrospective identification of chromatin signatures that bias cell state transitions. More specifically, under conditions of heterogeneous differentiation, one could group cells by their endpoint chromatin states, and then identify common, earlier, fate-determining chromatin signatures. Second, LagTag could be combined with imaging-based readout technologies to measure spatial proximities^48,49^ or molecular interactions^50^ between DNA loci and antibodies, potentially enabling spatiotemporal single-cell chromatin profiling, including past cell state information, directly within tissues.

LagTag has several limitations in its current form. First, the current study was performed on bulk populations, and a single-cell version remains to be developed. Second, m6A marks do not propagate and therefore dilute out over multiple rounds of cell division. Synthetic engineering of m6A replication mechanisms or the use of methylation dependent Dam mutants could help to address this issue^51,52^. Third, the current 16-hour m6A labeling window may exceed the cell cycle time of some fast-dividing cells, masking biological changes shorter than the labeling window. Fourth, non-specific adenine methyltransferases such as M.EcoGII can methylate RNA as well as DNA, potentially leading to cellular toxicity. As we showed, this issue could be overcome by further engineering or evolving the enzymatic properties of M.EcolGII^36^ to eliminate RNA methyltransferase activity while preserving DNA methylation. Despite these caveats, the temporal chromatin profiling approach we present here provides a versatile framework for resolving previously hidden cell state dynamics.

After posting the preprint describing this work, two manuscripts were posted, describing related approaches^53,54^. We provide a summary comparison of recent temporal chromatin profiling methods including these studies (**Table S3**). Collectively, this work should provide a foundation for applying chromatin recording to diverse challenges.

---

**Figure S1.**
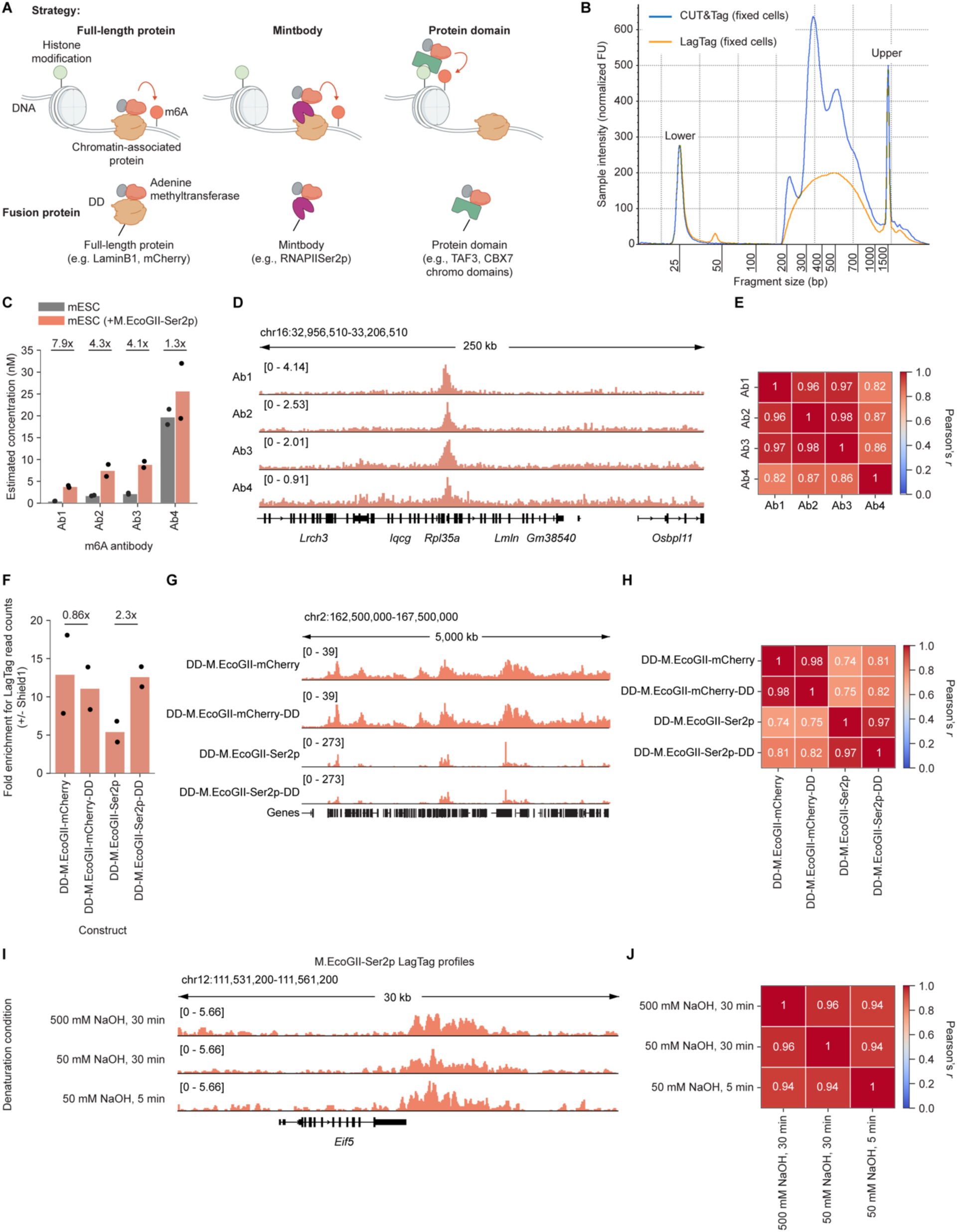
Optimization of CUT&Tag and LagTag for fixed cells. **A**, Schematic overview of different targeting strategies used for LagTag. **B**, Representative Tapestation profiles for LagTag and CUT&Tag in fixed cells. The profiles were obtained from RNAPIISer2p in mESCs. We note that the optimized condition of using proteinase K during the elution step, introduced by FFPE CUTAC^55^ for heavily crosslinked samples, also improves the yield of CUT&Tag and LagTag products from fixed cells by 1% formaldehyde directly on 96-well plates (see **Methods**). **C-E**, Comparison of different m6A antibodies (**Table S1**) with the LagTag assay in wild-type and M.EcoGII-Ser2p-mintbody-expressing mESCs. Ab1 was selected as a default m6A antibody based on its dynamic range. The yields of PCR products were estimated by the tapestation profiles within 200-1,000 bp in **C**. **F-H**, Comparison of different M.EcoGII fusion protein constructs, including different numbers of DD, with the LagTag assay in mESCs. The fold-enrichment with the addition of Shield-1 was calculated by comparing the read counts of samples mixed after PCR. **I**, **J**, Comparison of LagTag profiles across different denaturation conditions using M.EcoGII-Ser2p-mintbody-expressing mESCs. We note that the most harsh treatment condition of 500 mM NaOH for 30 minutes was used in the immunofluorescence assay in MadID^15^. Scale of LagTag profiles was normalized to coverage per million aligned reads in **D**, **G**, **I**. Correlation heatmaps of LagTag profiles were computed by using 10-kb bins across the genome in mESCs in **E**, **H**, **J**. n = 2 for experiments in **c**-**j**.

**Figure S2.**
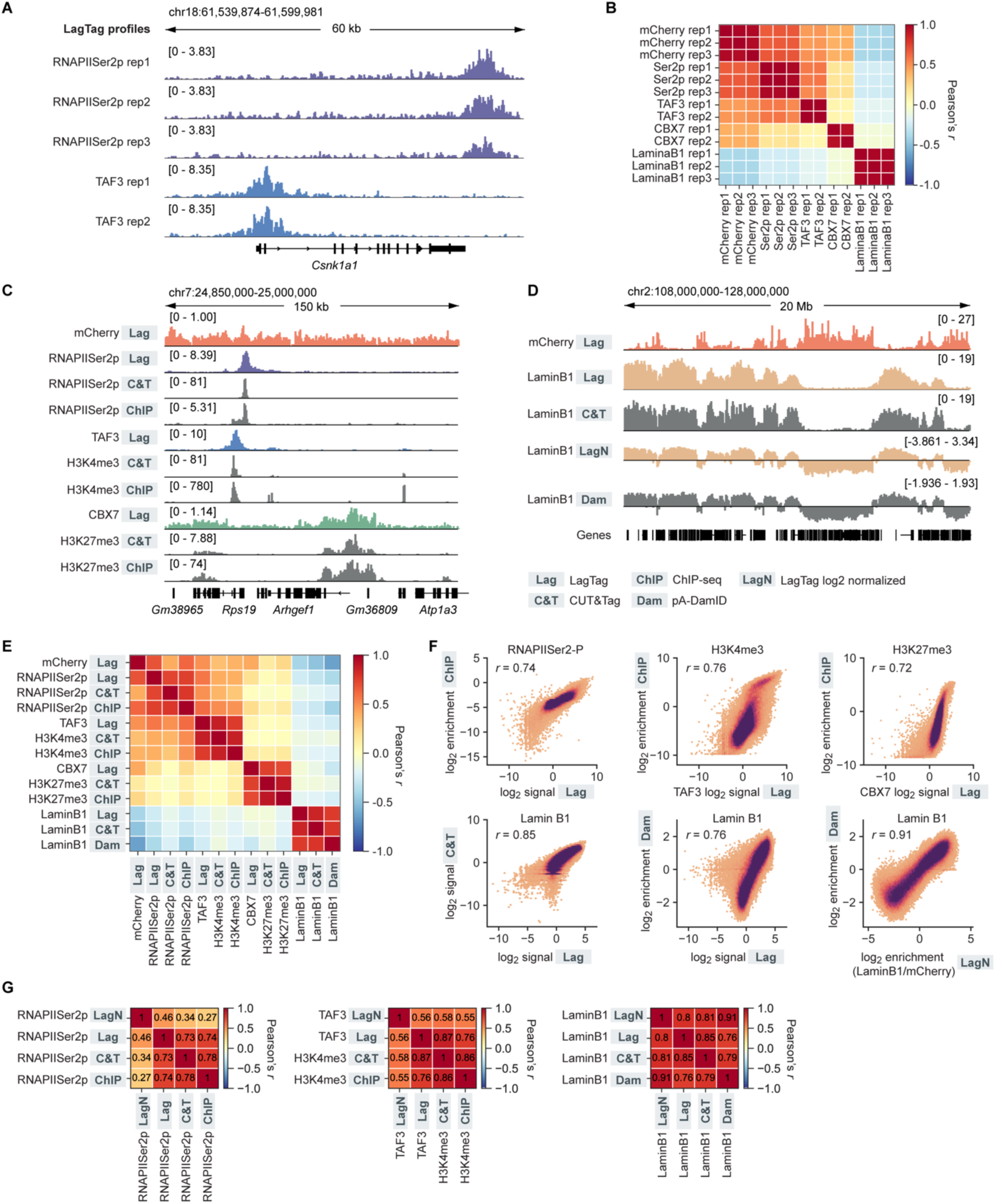
Validation and characterization of LagTag profiles in mESCs. **A**, **B**, Comparison of LagTag profiles between well replicates in mESCs. 10-kb bins were used for comparison in **B**. **C**, **D**, Genome browser tracks of LagTag, CUT&Tag, ChIP-seq^20,37^, and pA-DamID^33^. Scale of LagTag and CUT&Tag profiles was normalized to coverage per million aligned reads or normalized to log2 ratio between Lamin B1 and mCherry profiles (Lamin B1 LagTag normalized in **D**). The ChIP-seq profiles are input normalized and pA-DamID profiles are normalized by the Dam only control. **E**, Correlation heatmaps of chromatin profiles for different targets as well as different profiling methods using 10-kb bins across the genome in mESCs. **F**, Genome-wide comparison of chromatin profiles by different methods. r represents Pearson correlation coefficient shown in **E**. **G**, Correlation heatmap comparing LagTag profiles and log2(target/mCherry)-normalized LagTag profiles against orthogonal datasets. n = 2 for TAF3 and CBX7, and n = 3 for mCherry, RNAPIISer2p, and Lamin B1 in LagTag experiments; n = 2 for CUT&Tag experiments.

**Figure S3.**
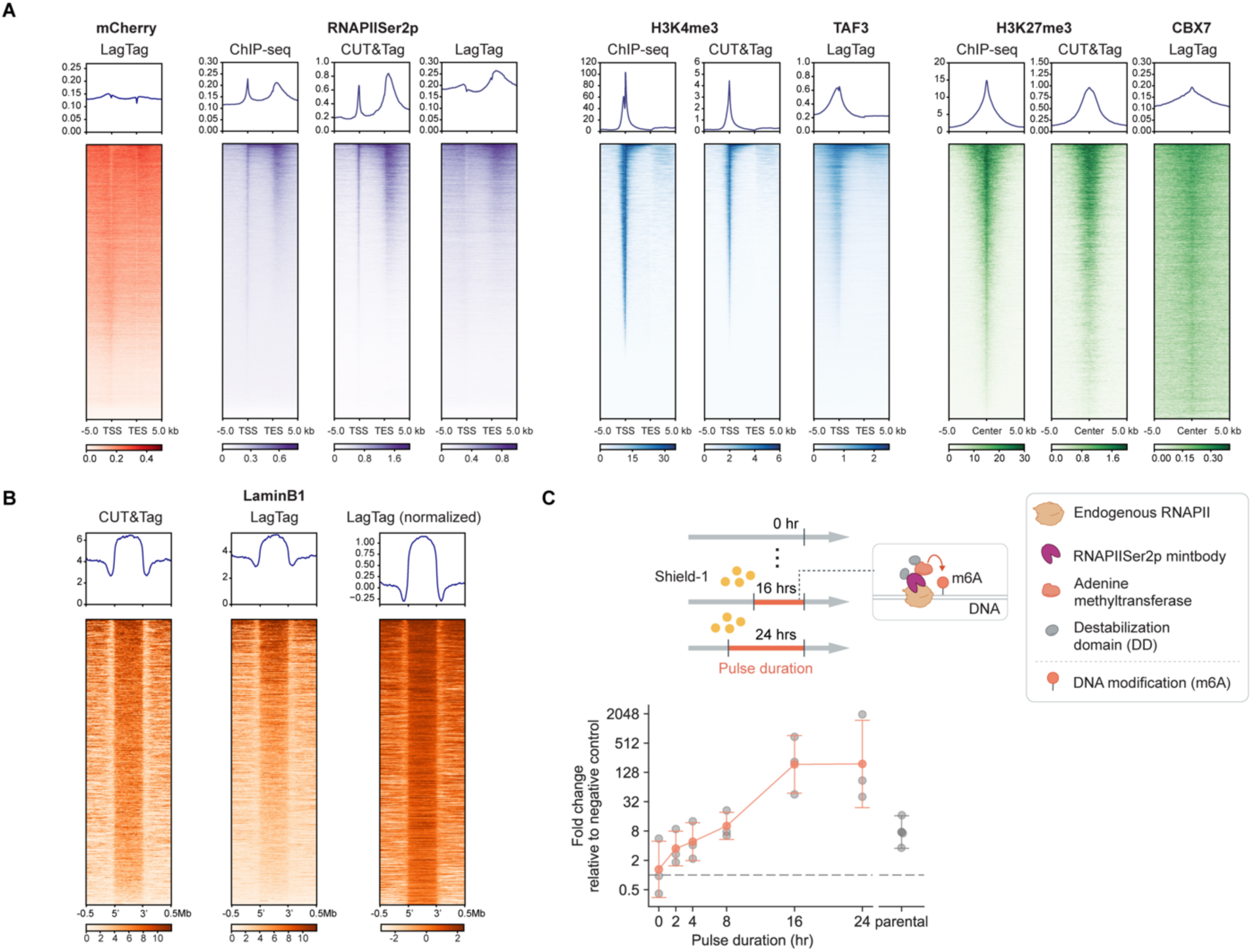
Additional Validation and characterization of LagTag profiles in mESCs. **A**, **B**, Signal enrichment plots for ChIP-seq^20,37^, CUT&Tag, and LagTag around TSS and TES across all protein-coding genes for mCherry, RNAPIISer2p, H3K4me3, or around called peaks from ChIP-seq^23,56^ for H3K27me3, or around LADs defined by DamID^57^ for Lamin B1 in mESCs. Scale of LagTag and CUT&Tag profiles was normalized to coverage per million aligned reads or normalized to log2 ratio between Lamin B1 and mCherry profiles (Lamin B1 LagTag normalized in **B**). n = 2 for TAF3 and CBX7, and n = 3 for mCherry, RNAPIISer2p, and Lamin B1 in LagTag experiments; n = 2 for CUT&Tag experiments. **C**, Comparison of LagTag yield as a function of Shield-1 pulse duration. Fold change was calculated relative to primer-only controls by qPCR (see **Methods**). We note that the m6A signal deposited by LagTag represents a temporal integral of chromatin states during the pulse duration, consistent with biophysical modeling of adenine methyltransferase kinetics^58^. The LagTag pulse duration is similar to or shorter than the other temporal chromatin methods (**Table S3**). The DD-M.EcoGII-Ser2p-mintbody-DD-expressing mESCs and parental mESCs (control) were used for this assay. Error bars represent mean ± standard deviation with individual replicate data points overlaid (n = 3).

**Figure S4.**
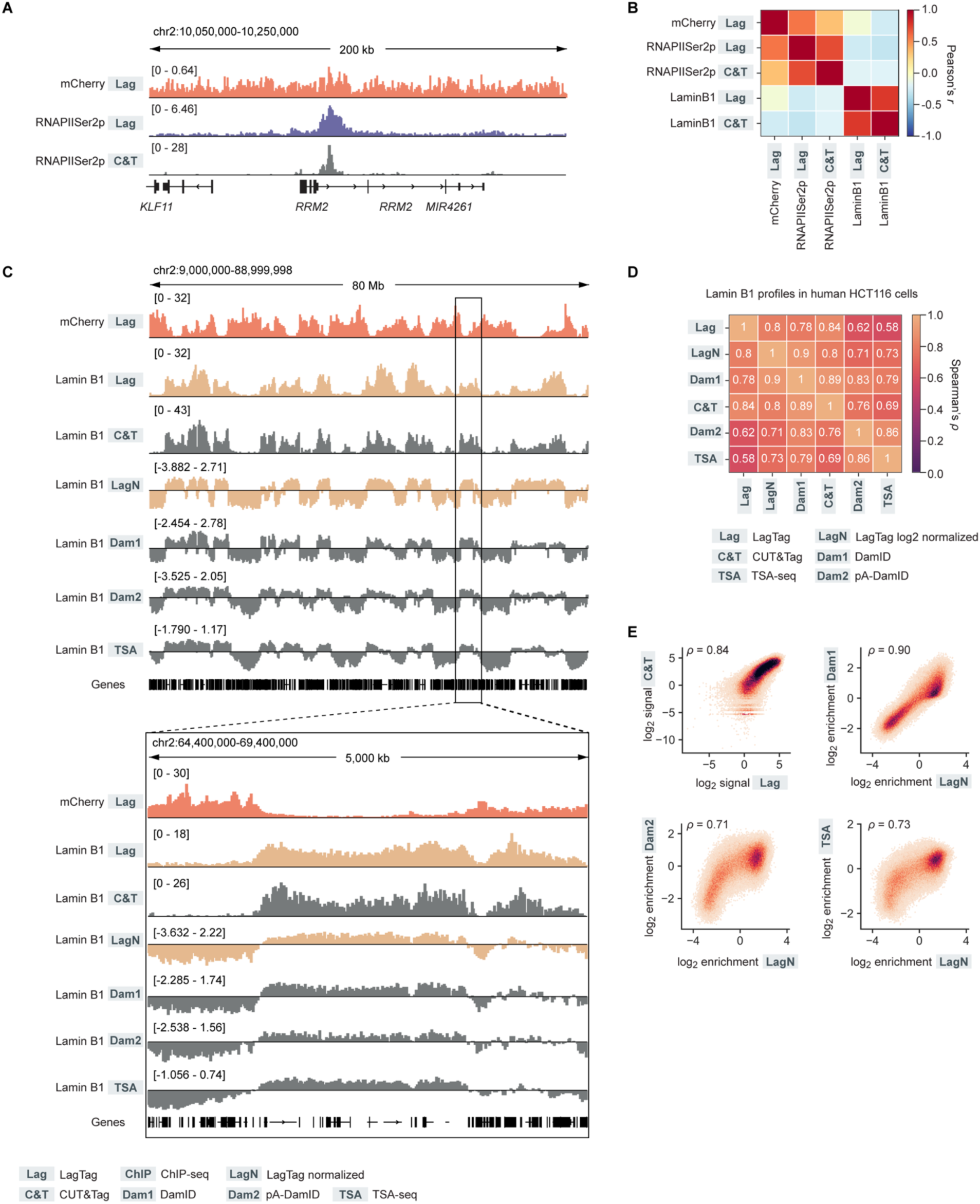
Validation and characterization of LagTag profiles in human HCT116 cells. **A**, Genome browser tracks of LagTag and CUT&Tag in human HCT116 cells. **B**, Correlation heatmaps of chromatin profiles for different targets as well as different profiling methods using 20-kb bins across the genome in HCT116 cells. **C**, Genome browser tracks of LagTag, CUT&Tag, DamID^59,60^, pA-DamID^61^, and TSA-seq^62^ in human HCT116 cells. Scale of LagTag and CUT&Tag profile was normalized to coverage per million aligned reads in **A**, **C,** or normalized to log2 ratio between Lamin B1 and mCherry profiles (LagTag normalized). The DamID and pA-DamID profiles are normalized by the Dam only control and the TSA-seq profiles are input normalized. **D**, **E**, Additional comparison of Lamin B1 profiles across different methods and normalization with commonly profiled 20-kb bins. n = 2 for CUT&Tag and LagTag experiments.

**Figure S5.**
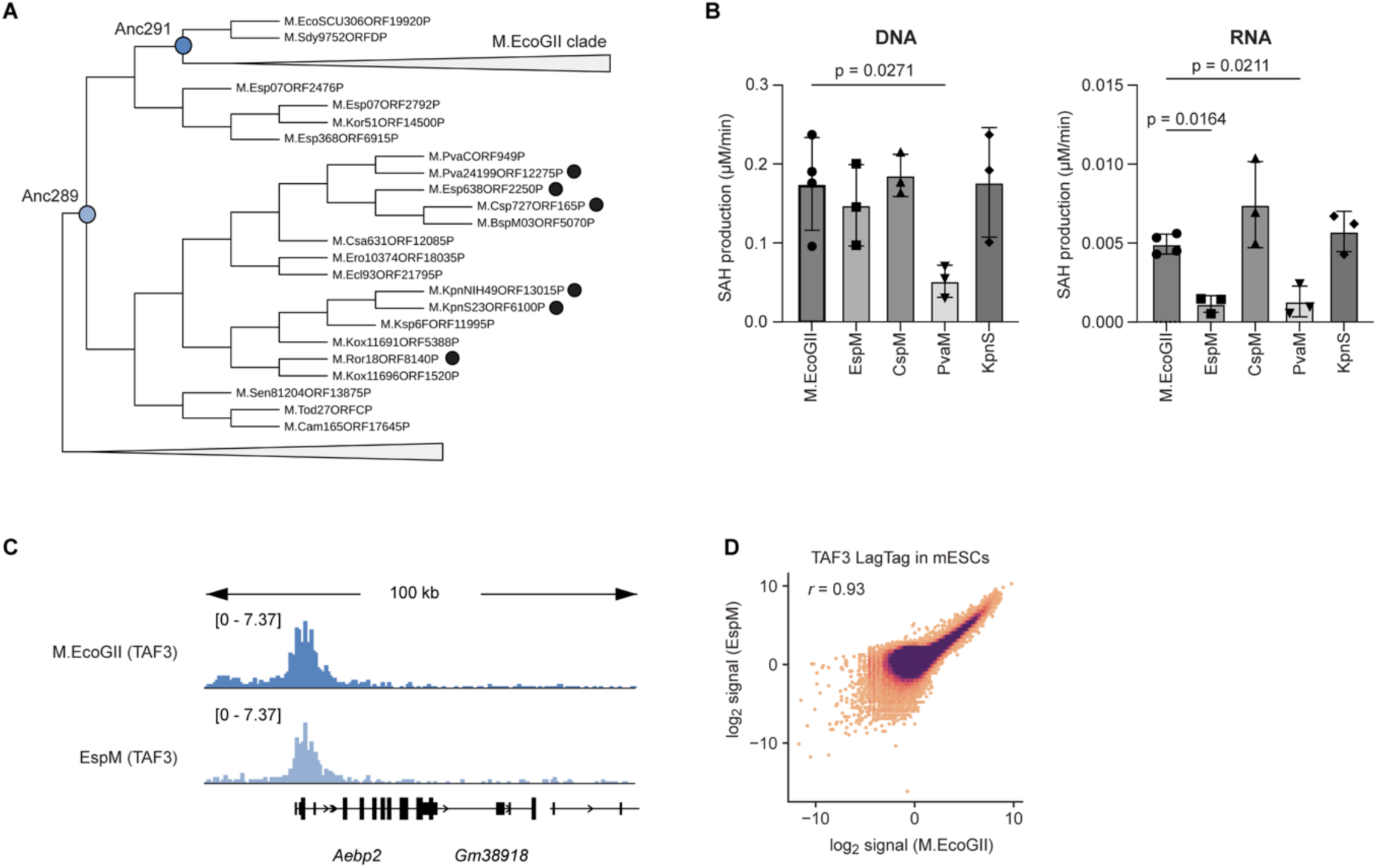
Comparison of m6A methylation levels and LagTag profiles between M.EcoGII variants. **A**, Identification and selection of M.EcoGII variants. The variants with black circles were selected for protein purification (see **Methods**). **B**, Comparison of in vitro methyltransferase activities on DNA (left) and RNA (right) across different M.EcoGII variants (see **Methods**). EspM showed lower RNA methyltransferase activity relative to M.EcoGII in vitro. n = 4 for M.EcoGII and n = 3 for other variants. **C**, Genome browser tracks of LagTag for TAF3 in mESCs by using fusion proteins with M.EcoGII or its variant EspM. **D**, Genome-wide comparison of LagTag profiles with M.EcoGII variants expressed in mESCs using 10-kb bins. r represents Pearson correlation coefficient. n = 2 for M.EcoGII and n = 3 for EspM in **C** and **D**.

**Figure S6.**
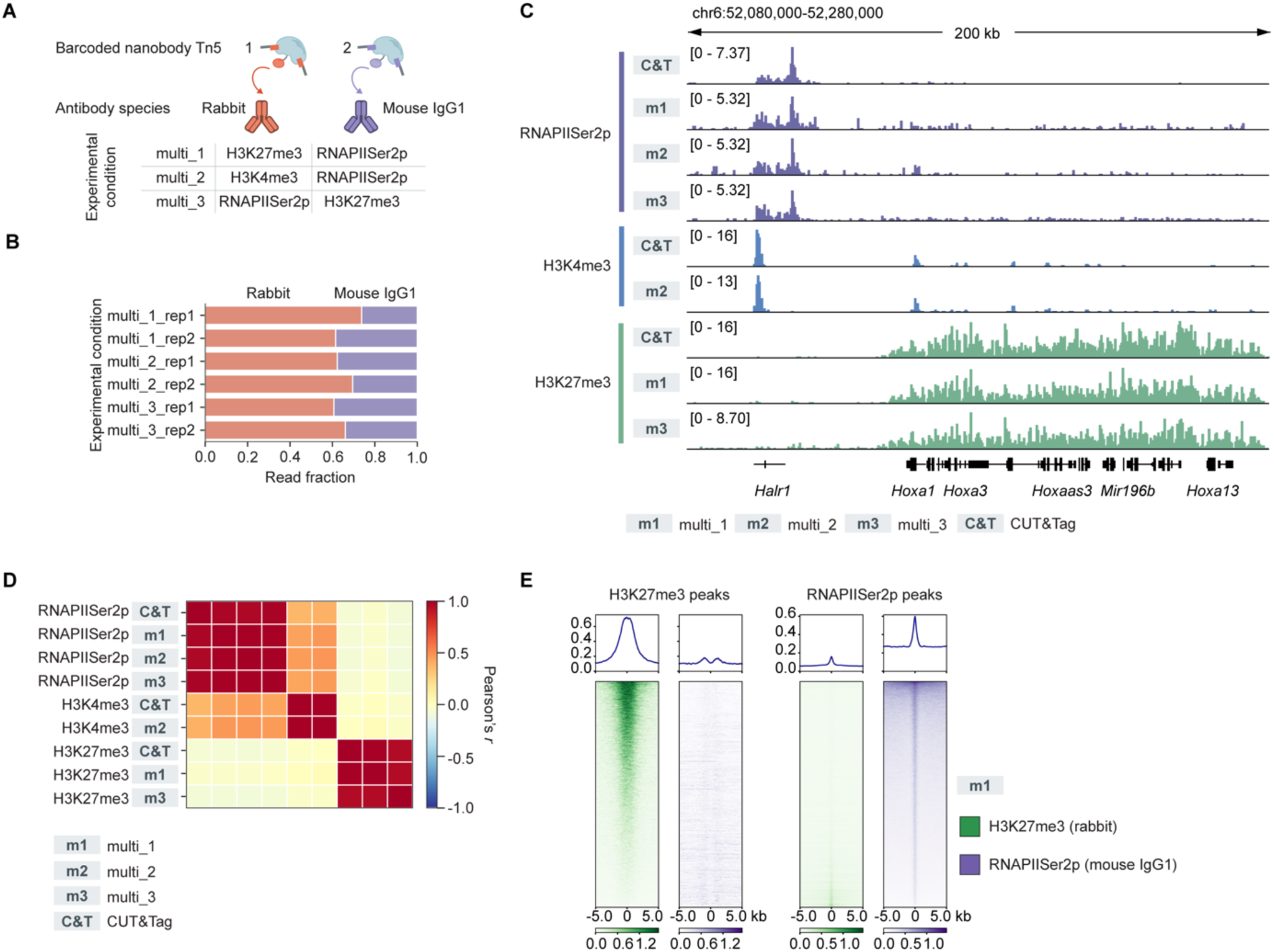
Validation of multifactorial CUT&Tag in mESCs. **A**, Representation of different antibody pairs used in multifactorial CUT&Tag in mESCs. The different antibodies were bound by antibody-species-specific nb-Tn5 with barcoded adapters. The assays were performed using fixed cells in 96-well plates. **B**, Comparison of read fraction from two target species after demultiplexing from each sample across conditions and replicates. The read fraction could be determined by the target abundance and antibody quality. The experimental condition and coloring of antibody species are shown in **A**. **C**, Representative genome browser tracks of CUT&Tag with non-barcoded adapters and multifactorial CUT&Tag with barcoded adapters. Scale of the profiles was normalized to coverage per million aligned reads. **D**, Correlation heatmaps of chromatin profiles for different targets using 10-kb bins across the genome. **E**, The comparison of enrichment of H3K27me3 (green) or RNAPIISer2p (purple) signal from multifactorial CUT&Tag at peaks provided by publicly available datasets (H3K27me3 ChIP-seq^23,56^ and RNAPIISer2p CUT&Tag^23^). n = 2 for experiments in **B**-**E**.

**Figure S7.**
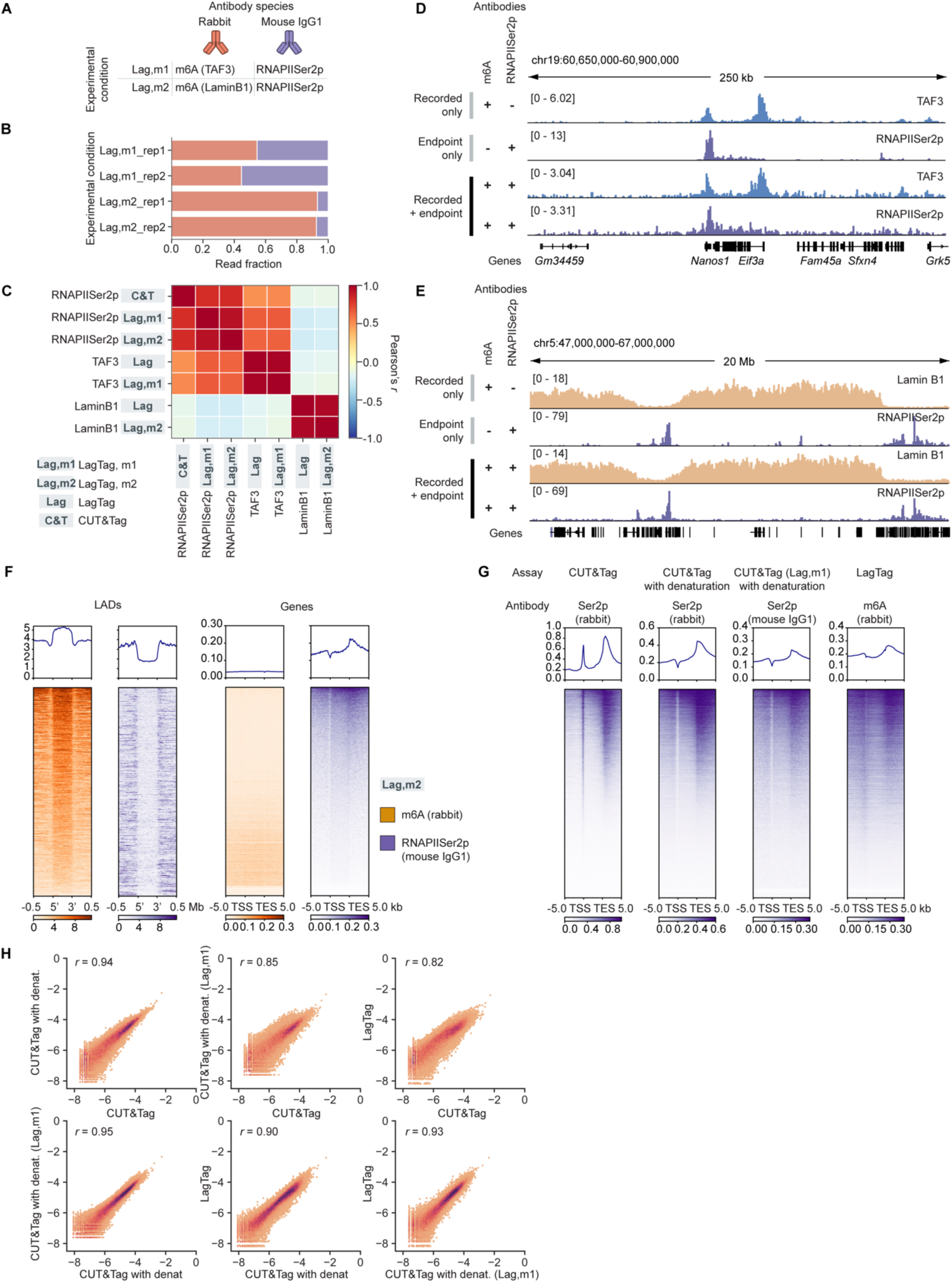
Validation of recorded and endpoint LagTag profiles. **A**, Representation of different antibody pairs used in LagTag assays in mESCs. Two different cell lines expressing M.EcoGII-TAF3 (reader domain) or M.EcoGII-Lamin B1 were used. **B**, Comparison of read fraction from two target species after demultiplexing from each sample across conditions and replicates. The experimental condition and coloring of antibody species are shown in **A**. **C**, Correlation heatmaps of chromatin profiles for different targets by LagTag with antibody pairs or individual assays using 10-kb bins across the genome. **D**, **E**, Representative genome browser tracks of LagTag profiles with barcoded adapters along with individual CUT&Tag and LagTag profiles with non-barcoded adapters in mESCs. Scale of the profiles was normalized to coverage per million aligned reads. 50-bp binning in **D** and 10-kb binning in **E** were used. **F**, Signal enrichment plots for LagTag condition 2 in **A** around LADs defined by DamID^57^ (left panels) and around TSS and TES across all protein-coding genes (right panels) in mESCs. Scale of the profiles was normalized to coverage per million aligned reads. **G**, Signal enrichment plots of RNAPIISer2p with CUT&Tag, CUT&Tag with denaturation of 50 mM NaOH for 5 minutes, CUT&Tag with denaturation of 50 mM NaOH for 5 minutes from experimental condition 1 in **A**, and LagTag with denaturation of 500 mM NaOH for 30 minutes (from left to right) around TSS and TES across all protein-coding genes in mESCs. We note that we observed the loss of peaks at TSS upon treatments with NaOH for CUT&Tag. **H**, Scatter plots evaluating the similarity of RNAPIISer2p profiles from different assays at the gene bodies across the genome in mESCs. r represents Pearson correlation coefficient. n = 2 or n =3 (Lamin B1 LagTag, RNAPIISer2p LagTag) for CUT&Tag and LagTag experiments in **B**-**E**.

**Figure S8.**
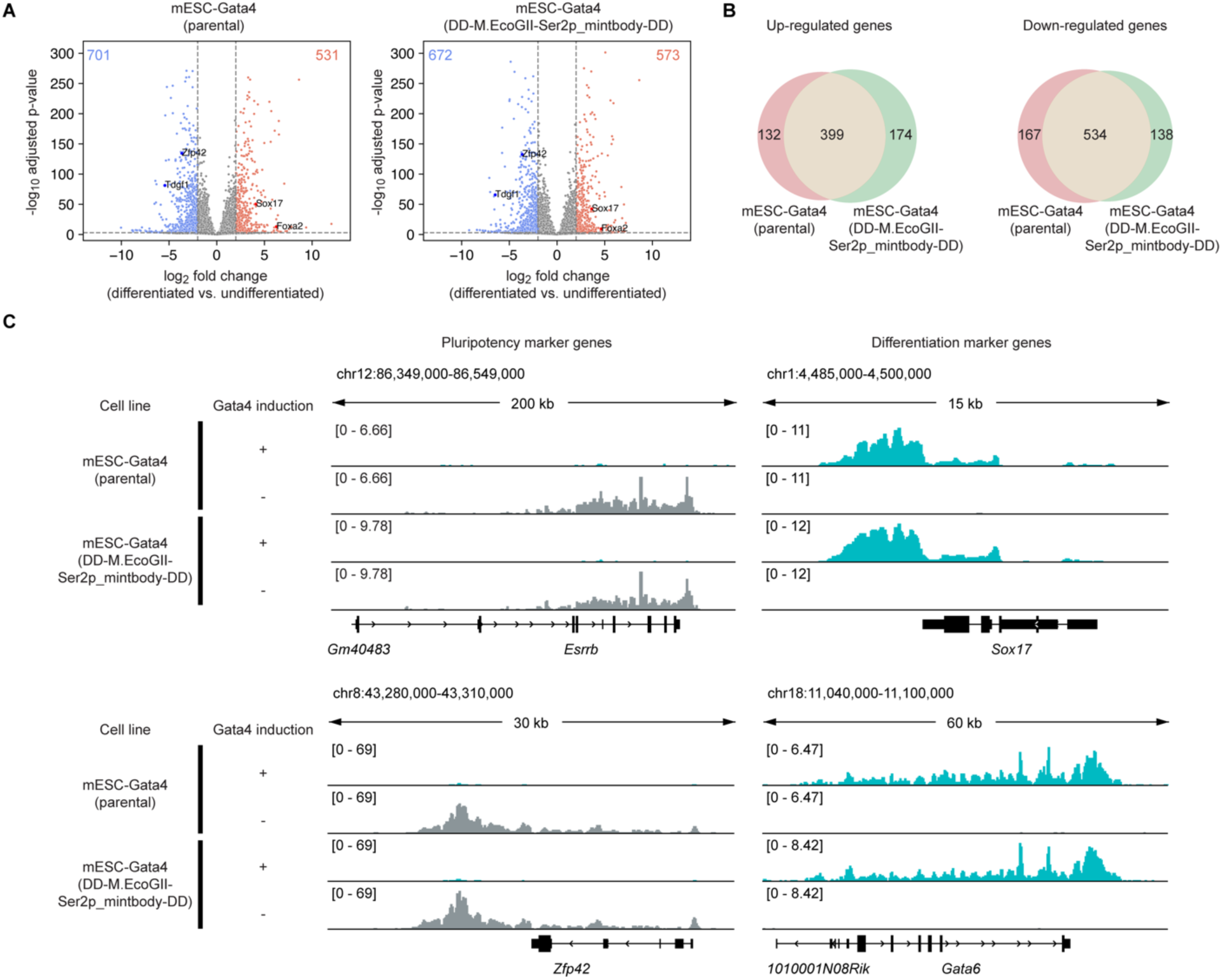
Characterization of mESC differentiation with an inducible Gata4 system. **A**, RNAPIISer2p CUT&Tag analysis comparing the signal enrichment over gene bodies before and after differentiation for 24 hours with the medium change in parental (left) or DD-M.EcoGII-Ser2p_mintbody-DD engineered (right) mESC-Gata4 cells. Single clones were used for these cell lines. Marker genes used in Figure 2D**, E** are highlighted with individual gene names. **B**, Up-regulated (left) and down-regulated (right) genes during the differentiation defined by RNAPIISer2p CUT&Tag in the mESC-Gata4 lines (see **Methods**). **c**, Representative genome browser tracks of RNAPIISer2p CUT&Tag comparing the undifferentiated (gray) and differentiated (cyan) mESC-Gata4 cells with marker genes. n = 3 for CUT&Tag experiments.

**Figure S9.**
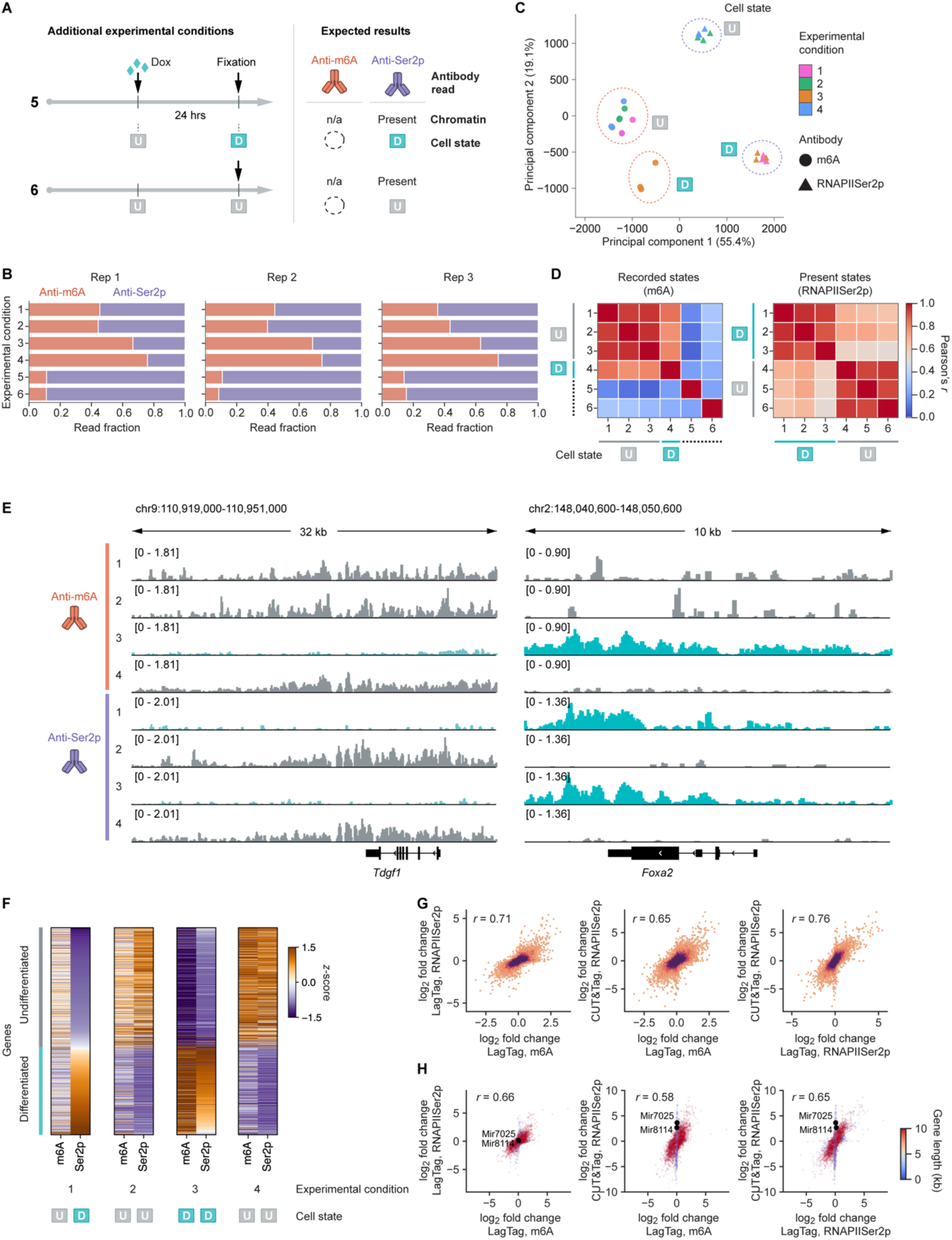
Validation and characterization of LagTag profiles during mESC differentiation. **A**, Additional schematic representation of LagTag experimental conditions with mESC differentiation along with expected results, corresponding to the endpoint chromatin states in each condition. The chromatin recording was not performed in these control conditions, while anti-m6A antibody was incubated with the samples. **B**, Comparison of read fraction from two target species after demultiplexing from each sample across conditions and replicates. **C**, Principal component analysis of CPM-normalized chromatin profiles (10-kb bins) colored by different experimental conditions along with the recorded and endpoint cell states. **D**, Correlation heatmaps of chromatin profiles from different experimental conditions with LagTag using 10-kb bins across the genome, along with the recorded (left) and endpoint (right) cell states. **E**, Genome browser tracks of LagTag samples across experimental conditions as well as two antibodies. The tracks are colored based on the recorded (m6A tracks) or endpoint (RNAPIISer2p tracks) cell states. **F**, Heat maps of z-score normalized LagTag profiles over genes differentially enriched with RNAPIISer2p (n = 933 genes) between undifferentiated and differentiated cell states, as determined by independent CUT&Tag experiments (n = 3). Z-score normalization was performed across conditions for each antibody (either m6A or RNAPIISer2p). Genes were sorted by RNAPIISer2p z-score enrichment in experimental condition 1. The same z-score normalized LagTag profiles, clustered by experimental conditions and genes, are shown in Figure 2F**. G**, **H**, Fold change in RNAPIISer2p signal enrichment over gene bodies during differentiation measured by LagTag (m6A or RNAPIISer2p antibody in condition 3 vs condition 4) or CUT&Tag (RNAPIISer2p only in condition 5 vs 6) in engineered mESC-Gata4 cells for coding genes (**G**) and all genes (**H**). Fold changes for some short genes were not captured by LagTag with NaOH denaturation compared with conventional CUT&Tag, potentially limiting the detection of a subset of genes by LagTag. Two representative short genes (Mir7025, Mir8114) are highlighted. n = 3 for LagTag and CUT&Tag experiments in **B**-**H**.

**Figure S10.**
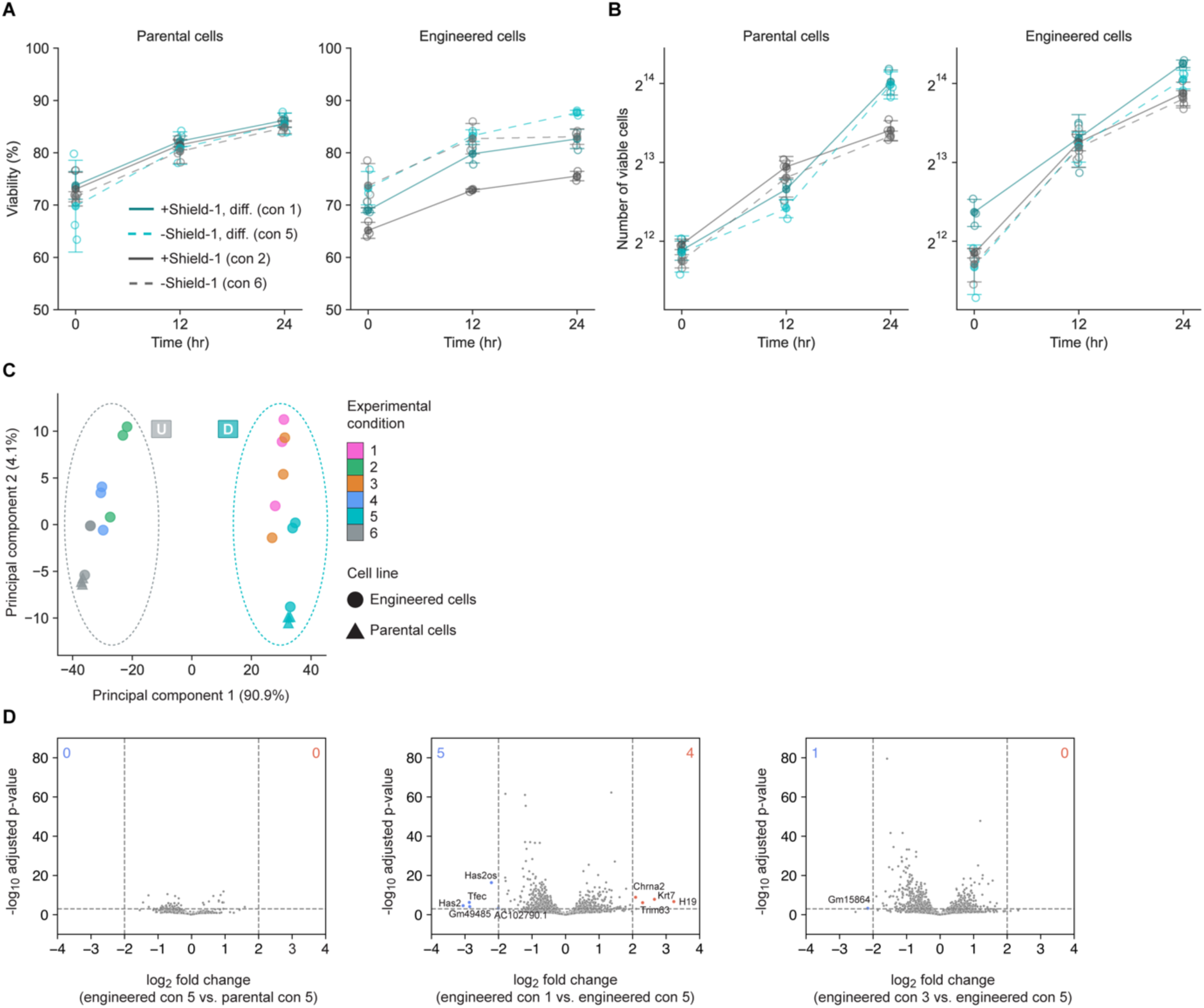
Evaluation of LagTag recording effects on cell physiology during mESC differentiation. **A**, **B**, Comparison of viability (**A**) and number of viable cells (**B**) with or without prior Shield-1 induction for 16 hours in undifferentiated and differentiated cells (n = 3). **C**, Principal component analysis of RNAPIISer2p CUT&Tag profiles across different experimental conditions in engineered and parental mESC-Gata4 cells using the 500 most variable genes. **D**, RNAPIISer2p CUT&Tag analysis comparing signal enrichment over gene bodies between conditions in parental or engineered mESC-Gata4 cells. The DD-M.EcoGII-Ser2p_mintbody-DD engineered and parental mESC-Gata4 cells were used in **A**-**D**. n =3 for CUT&Tag experiments in **C** and **D**.

## Acknowledgements

We thank J. Vielmetter and A. Lam at the Caltech Protein Expression Center in the Beckman Institute for protein purification services for purifying nb-Tn5 enzymes; I.-M. Strazhnik for illustrations and formatting the figures; M. Zernicka-Goetz for providing the mESC-Gata4 cell line; B. Emert, J. Parres-Gold, I. N. Goronzy for help with the analysis pipeline setup; L. Santat, J. Leonardo, and R. Malinowski for administrative support. This research was supported by the National Institutes of Health (award numbers U01DK127420, R01AG080982 and R01DK143671) and by the Paul G. Allen Frontiers Group (the Allen Discovery Center Award UWSC13408). The content is solely the responsibility of the authors and does not necessarily represent the official views of the National Institutes of Health. M.B.E. is a Howard Hughes Medical Institute Investigator. Y.T. was supported by the Riva Foundation Fellowship.

## Author contributions

Y.T. and M.B.E. conceived and designed the study. Y.T., J.M.L., R.H. prepared the engineered cell lines. Y.T. performed sequencing experiments. Y.T., J.A.L., D.M.C. performed sequencing data processing and analysis. A.A.P., M.R.B., M.G. assisted with or offered guidance regarding sequencing experiments and analysis. Y.O. and P.L. designed and performed in vitro methyltransferase experiments and analysis. Y.T. and J.M.L. performed validation experiments and analysis. M.B.E. directed and supervised the study. Y.T., J.A.L., and M.B.E. wrote the manuscript with input from all authors.

## Competing interests

Patent applications related to this work have been filed by the California Institute of Technology (Y.T. and M.B.E.). M.B.E. is a scientific advisory board member or consultant at TeraCyte, Plasmidsaurus, Asymptote Genetic Medicines, and Spatial Genomics. The other authors declare no competing interests.

**Materials & Correspondence** should be addressed to Michael Elowitz.

## Data availability

All sequencing datasets generated in this study are available at NCBI GEO (accession GSE305653) as of the date of publication. Additional processed data from this study are available at CaltechDATA (https://doi.org/10.22002/abcrz-g3683). Publicly available datasets used in this study (ENCSR212KGS, ENCSR059MBO, GSE195840, GSE171554, GSE181693, 4DNESWB729QB, 4DNES24XA7U8, 4DNESTE8NJOE, and https://zenodo.org/records/5507375) are detailed in the **Methods**.

## Code availability

Publicly available software and packages were used in this study as indicated in **Methods**. The custom written code used in this study is available at CaltechDATA (https://doi.org/10.22002/abcrz-g3683).

## Methods

### Cell lines

E14 mouse embryonic stem cells (E14Tg2a.4) were maintained on 0.1% gelatin (Sigma-Aldrich G1393) coated plates under the serum/LIF condition in DMEM (Gibco 11960044) supplemented with 15% ES-grade fetal bovine serum (FBS, Gibco 16141061), 1,000 units/mL leukemia inhibitory factor (LIF) (Sigma-Aldrich ESG1106), 1× non-essential amino acids (Gibco 11140050), 1 mM sodium pyruvate (Gibco 11360070), 55 μM 2-mercaptoethanol (Gibco 21985023), and 1× Penicillin-Streptomycin-Glutamine (Gibco 10378016). mESC-Gata4 cells^22^ were kindly provided by Magdalena Zernicka-Goetz and maintained on 0.1% gelatin coated plates under the serum/LIF/2i condition by supplementing the serum/LIF medium with 1 μM PD0325901 (Sigma-Aldrich 444966) and 3 μM CHIR99021 (Sigma-Aldrich SML1046). The mESC differentiation to endodermal cells was performed with 1 μg/mL Dox induction and LIF withdrawal for 24 hours. The human HCT116 (ATCC CCL-247) cells were maintained in McCoy’s 5A (modified) medium (Gibco 16600082) supplemented with 10% FBS (Avantor 97068-085) and 1× Penicillin-Streptomycin (Gibco 15140122). The cell lines were not routinely tested for mycoplasma contamination. All cell lines were cultured at 37°C with 5% CO2.

### LagTag construction design

Plasmids were constructed in a PiggyBac vector (System Biosciences) with a puromycin resistance gene using standard cloning methods. The adenine methyltransferase fusion genes are expressed under the PGK promoter and mutated Kozak sequence, TGATAT, for lower translation efficiency^29^. The DD-linker-M.EcoGII was cloned from pRetroX-PTuner DD-linker-M.EcoGII (Addgene 122082) and DD-linker-M.EcoGII-LaminB1 (LMNB1) was cloned from pRetroX-PTuner DD-linker-M.EcoGII-v5-LaminB1 (Addgene 122083)^15^. The Ser2p-mintbody^31,63^, TAF3 (three chromo domains)^28^, CBX7 (two chromo domains)^27^, and EspM were cloned from gBlocks (Integrated DNA technologies) to make the similar constructs. The DD was further amplified by PCR and cloned to make dual DD constructs.

### Generation of stable LagTag cell lines

To make stable cell lines with adenine methyltransferase fusion constructs, the cells were transfected with 50 µg of the PiggyBack vectors on a 24-well plate using FuGENE HD transfection reagent (Promega E2311). The polyclonal cells were selected using 1 μg/mL puromycin and single clones for mESCs were manually isolated for some of the experiments. The adenine methyltransferase fusion proteins were stabilized with 1 μM Shield-1 (Takara Bio 632189) for 16 hours.

### Exploration of M.EcoGII homologs

Twenty-six M.EcoGII homologs between anc291 and anc289 nodes were selected from the maximal likelihood tree constructed previously^36^. Two unaligned sequences were excluded from the multiple sequence alignment (MSA). 19 of 24 sequences were selected based on conservation of residue Q179, which is a key residue for enhanced RNA methylation activity. One unaligned node was excluded using a sequence similarity network (SSN) generated from the sequences. Sequence similarity was calculated by Ident and Sim^64^ and similarity The amino acid set was used for similarity calculation: [GAVLI, FYW, CM, ST, KRH, DENQ, P]. Finally, 6 sequences were selected by sequence similarity less than 90% to two ancestral M.EcoGII variants: Anc291 and Anc289.

A sequence encoding a C-terminal His6 tag with a linker sequence (N-GGAAALGHHHHHH-C) was added to each candidate gene, and the resulting DNA sequences were synthesized and cloned into the pET-28a (+) plasmid using seamless cloning method. In-frame cloning into the pET-28a (+) vector was confirmed by colony PCR and DNA sequencing.

Plasmids containing candidates were transformed into E. coli strain SHuffle T7 (C3026J, NEB) and plated on LB agar containing 30 μg/mL kanamycin and 2% glucose, incubating at 37°C overnight. Colonies of CspM were inoculated into 10 mL of LB media with 30 μg/ml kanamycin and 2% glucose and incubated at 37°C with shaking at 220 rpm overnight. The preculture was transferred into 500 mL of LB media with 30 μg/mL kanamycin and 1% glucose, and the E. coli cells were grown at 37°C with shaking at 220 rpm until the OD600 reached 0.5–0.8. Colonies of EspM, PvaM, or KpnNIH were directly inoculated into 750 mL LB media with 30 μg/mL kanamycin and 1% glucose without preculture, and the E. coli cells were grown at 37°C with shaking at 220 rpm until the OD600 reached 0.5–0.8. Protein expression of DNA selective MTase candidates was induced by 1 mM isopropyl β-d-1-thiogalactopyranoside (IPTG) (EI07011, Biosynth) for 18 hours at 18°C. Cells were harvested and stored at −20°C before protein purification.

Cells were resuspended with lysis buffer (20 mM HEPES (pH 7.0), 250 mM NaCl, and 10 mM imidazole) and lysed by sonication. The lysis supernatant was obtained by centrifugation at 10,000 × g, 4°C for 30 min. M.EcoGII homologs were purified by affinity chromatography in an open column, eluting with 20 mM HEPES (pH 7.0), 250 mM NaCl and 100 mM imidazole at room temperature. Each fraction was analyzed by SDS–PAGE. Fractions enriched in the target protein were pooled, filtered through a 0.22 μm filter and concentrated in an Amicon Ultra 30 kDa MWCO centrifugal concentration device (UFC9030, Merck) with working buffer (20 mM HEPES (pH 7.0), 250 mM NaCl). The concentrated protein samples (>1 mg/mL protein and <1 mM imidazole) were mixed with glycerol to reach 10% (v/v) concentration, quickly frozen by liquid nitrogen and stored at −80°C.

### In vitro methyltransferase activity measurement

The DNA substrate encoding M.EcoGII (3569 bp, containing 1819 adenine bases) was amplified by PCR and purified using a NucleoSpin Gel and PCR Clean-up Kit (740609, MACHEREY-NAGEL) with Milli-Q water. The RNA substrate encoding the firefly luciferase gene (1872 nt, 544 adenine bases) was prepared by in vitro transcription using HiScribe T7 High Yield RNA Synthesis Kit (E2040S, NEB) with 1 unit RNase Inhibitor and purified in ribonuclease (RNase)-free water using an RNA Clean & Concentrator-5 kit (R1015, Zymo Research). In vitro methylation experiments were performed for 10 minutes (DNA) or 1 hour (RNA) at 37°C in 8 μL reaction mixtures containing 1× CutSmart buffer with 20 nM DNA or 50 nM RNA with 1 unit RNase Inhibitor (M0314S, NEB), 100 nM methyltransferase and 20 μM SAM. The methylation reaction in the 8 μL reaction was quenched by heating at 80°C for 1 min at each timepoint. The methyltransferase reaction product S-adenosylhomocysteine was quantified using MTase-Glo kit (V7601, Promega) following the manufacturer’s protocol^65^. To normalize the S-adenosyl homocysteine (SAH) production rate in DNA/RNA methylation, a calibration curve for SAH was constructed in two independent experiments using from 0 to 5 μM SAH according to the manufacturer’s protocol of the MTase-Glo kit. The signal was detected using a Tecan Spark microplate reader (Tecan Japan Co., Ltd).

### Nanobody-Tn5 transposome preparation

The pTXB1-nbOcIgG-Tn5 (Addgene 184285) and pTXB1-nbMmIgG1-Tn5 (Addgene 184287) plasmids^25^ were purchased from Addgene. The nb-Tn5 transposase production was performed as previously described^25^ with modifications. The plasmid was transformed into ArcticExpress (DE3) Competent Cells (Agilent Technologies 230192), and 1 liter of terrific broth culture was grown at 37 °C until the OD600 reached 0.6. The nb-Tn5 expression was then induced with 0.25 mM isopropyl-ß-D-thiogalactopyranoside (IPTG) at 22 °C for 18 hours. The cells were then pelleted and stored at -80 °C until use. For protein purification, the cells were lysed by sonication in 100 mL of HEGX buffer, consisting of 20 mM HEPES-KOH pH 7.5, 1 M NaCl, 1 mM EDTA, 10% glycerol, 0.2% Triton X-100, and a protease inhibitor cocktail (Roche 04693132001). The lysate was then pelleted at 30,000g at 4 °C for 20 minutes. The supernatant was transferred to a new tube, and 10% polyethylenimine (Sigma-Aldrich P3143) was added to the bacterial extract, followed by gentle mixing and centrifugation at 30,000g at 4 °C for 30 minutes. The supernatant was then loaded on an Econo-Column column (Bio-Rad Laboratories 7371022) filled with 10 mL of chitin resin (NEB S6651L). The column was washed with 50 mL of HEGX buffer, and then incubated with 15 mL of HEGX buffer supplemented with 100 mM DTT at 4 °C for 48 hours to cleave the nb-Tn5 from the intein tag. The nb-Tn5 was eluted directly into one 10 kDa MWCO Amicon Ultra Centrifugal Filter (Millipore UFC9010) by the addition of 10 mL of HEGX buffer. The protein was spin concentrated in the Amicon Ultra Centrifugal Filter using 15 mL of 2× dialysis buffer, consisting of 100 mM HEPES-KOH pH 7.2, 0.2 M NaCl, 0.2 mM EDTA, 2 mM DTT, 20% glycerol, and concentrated to 1 mL by centrifugation at 5,000g. The protein concentrate was transferred to a new tube and mixed to make a final concentration of 50% glycerol. The nb-Tn5 aliquots were stored at -80 °C.

The nb-Tn5 adapter transposomes were assembled as previously described^26^. Briefly, 100 µM equimolar mixture of two adaptor oligonucleotides (**Table S2**) in an annealing buffer, consisting of 10 mM Tris-HCl pH 8.0 (Invitrogen 15568025), 50 mM NaCl (Invitrogen AM9759), 1 mM EDTA pH 8.0 (Invitrogen AM9260G) in DNase/RNase-free distilled water (Invitrogen 1097701), was denatured on a thermocycler for 5 minutes at 95°C and cooled down at room temperature for more than 45 minutes for annealing. The 100 µM equimolar mixtures (50 µL of each) were mixed with 500 µL of 6.8 µM nb-Tn5 in the 2× dialysis buffer and incubated for 1 hour at room temperature. The transposomes were mixed with 100% glycerol to make a final concentration of 50% glycerol and stored at -20 °C until use.

### On-plate LagTag and CUT&Tag

LagTag and CUT&Tag experiments were performed using adherent cells directly fixed on the 96-well plate. The conventional CUT&Tag protocol^66^ was optimized for this implementation based on the recent chromatin profiling technologies^55,67,68^ and by incorporating the nanobody-based transposome^25,26^. E14 and mESC-Gata4 cells (6,000-12,000 cells) were seeded on human laminin (BioLamina LN511) coated 96-well plates, and HCT116 cells (10,000 cells) were seeded on collagen (Sigma-Aldrich C8919) coated ibiTreat 96-well plates (ibidi 89606). Cells were washed with 100 μL of DPBS (Gibco 14190144) once and fixed with 50 μL of freshly-prepared 1% formaldehyde (Thermo Scientific 28908) buffer in 1× PBS (Invitrogen AM9624) at room temperature for 5 minutes. The fixed cells were washed twice with 100 μL of DPBS and used on the same day. The DNase/RNase-free distilled water was used throughout the experiments. The cells were permeabilized with 50 μL of 0.5% Triton-X (Sigma-Aldrich 93443) in 1× PBS at room temperature for 20 minutes and washed once with 50 μL of 1× PBS at room temperature. For LagTag, to remove endogenous m6A RNA modifications, cells were incubated with 50 μL of 5% RNaseA/T1 Mix (Thermo Fisher EN0551) in 1× PBS at 37°C for 1 hour. Then cells were denatured^15^ with 30 μL of a denaturation buffer, consisting of 1.5 M NaCl (Invitrogen AM9759) and 500 mM NaOH (Teknova H0224), at room temperature for 30 minutes, followed by three washes with 50 μL of a neutralization buffer, consisting of 500 mM Tris-HCl pH 7.5 (Invitrogen 15567027) and 2.5 M NaCl, at room temperature. For LagTag with multi-antibody samples, cells were denatured with a milder condition of 150 mM NaCl and 50 mM NaOH at room temperature for 5 minutes, followed by three washes with a neutralization buffer (50 mM Tris-HCl pH 7.5 and 250 mM NaCl). The samples were further washed with 50 μL of 1× PBS at room temperature three times. Those RNase and denaturation steps were required specifically for LagTag and were omitted in CUT&Tag experiments unless otherwise specified. The samples were then washed once with 1× PBS and blocked with 20 μL of a blocking solution consisting of 1× PBS, 10 mg/mL UltraPure BSA (Invitrogen AM2616), 0.05% Triton-X at room temperature for 5-15 minutes. Then the cells were incubated with 30 μL of 100-fold diluted primary antibodies in the blocking solution at 4°C overnight. Primary antibodies used in this study are listed in **Table S1**. On the following day, the samples were washed twice with 50 μL of wash 300 buffer, consisting of 20 mM HEPES pH 7.4 (Teknova H1030), 300 mM NaCl, and 0.05% Triton-X, and then incubated with 30 μL of wash 300 uL buffer supplemented with 1.5-3 μL of nb-Tn5 transposome (in-house) and 0.1% BSA at room temperature for 1 hour with a gentle shaking of 300 rpm. For multi-antibody LagTag or CUT&Tag experiments with barcoded adapters, cells were incubated at 4°C overnight in 30 μL of wash 300 buffer supplemented with 2 μL of each barcoded nb-Tn5 transposome and 0.1% BSA. The samples were then washed twice with 50 μL of wash 300 buffer. The samples were then incubated with 50 μL of tagmentation buffer (10 mM MgCl_2_ (Invitrogen AM9530G) and 0.1% BSA in the wash 300 buffer) at 37°C for 1 hour. We note that the plates were sealed with adhesive PCR plate sealing films (Bio-Rad Laboratories MSB1001B) to prevent evaporation and placed in the 37°C incubator without stacking multiple plates to allow homogeneous heating on the plate surface. Then the samples were washed once with 50 μL of a TAPS buffer, consisting of 10 mM TAPS buffer pH 8.5 (Boston BioProducts BB-2375) and 0.2 mM EDTA. Then 20 μL of SDS release buffer, consisting of 10 mM TAPS buffer pH 8.5, 0.2% SDS (Invitrogen 24730020), and 5% thermolabile proteinase K (NEB P8111S), was added to each well, followed by an incubation at 37°C for 30 minutes and then at 62°C for 30 minutes. Similarly to the tagmentation step, plates were covered by the seal and each plate was placed on the surface of the incubator. The use of thermolabile proteinase K in the release buffer was recently introduced for heavily crosslinked formalin-fixed paraffin-embedded samples^55^, but we noticed it is applicable to recover longer fragments from crosslinked samples on the plate by 1% formaldehyde. The SDS release buffer was then mixed with 12 μL of 10% Triton-X for neutralization and used as a PCR input. The sequencing libraries were then prepared by adding 21 μL of the input, 2 μL of i5 and i7 primers (Integrated DNA technologies, 10 μM stock each) (**Table S2**), and 25 μL of NEBNext High-Fidelity 2X PCR Master Mix (NEB M0541L) to each PCR tube, followed by PCR amplifications: 5 min at 58°C, 5 min at 72°C, and 45 sec at 98°C followed by 13-15 cycles of 15 sec at 98°C and 10 sec at 60°C, with a final extension at 72°C for 1 min and a hold at 4°C. The amplified libraries were purified using 1.3× magnetic beads (Omega Bio-Tek M1378) according to the manufacturer’s protocol.

### Sequencing

After the pooled libraries were purified by E-Gel EX Agarose Gels, 2% (Invitrogen G402022) and spin column (Zymo D4001, D4014), the samples were quantified using the Qubit dsDNA High Sensitivity Assay (Invitrogen Q32854) and the Agilent High Sensitivity D1000 DNA ScreenTape Assay (Agilent Technologies 5067-5603, 5067-5584). Then the paired-end sequencing of the libraries was performed on the AVITI using the AVITI 2x150 Sequencing Kit Cloudbreak FS (Element Biosciences 860-00011, 860-00013) with Cloudbreak FS PhiX Control, 3rd Party (Element Biosciences 830-00023) according to the manufacturer’s protocol with a 2% PhiX spike-in. For the adapter-barcoded samples, custom primers^23^ (Integrated DNA technologies) were used for Read 1, Read 2, and Index 1 (**Table S2**) with Custom Primer Set Cloudbreak Freestyle (Element Biosciences 820-00025) and libraries were mixed with a 30% PhiX spike-in.

### Quantification of m6A deposition by qPCR

Tagmentation efficiency, reflecting the extent of m6A deposition, was assessed by qPCR as previously described for library quantification for ATAC-seq^69^ and CUT&Tag^70^. Briefly, 1 μL of purified LagTag amplicon was used as input for a qPCR assay using the corresponding i5 and i7 index primers at 250 nM with iQ SYBR Green Supermix (Bio-Rad) in a total reaction volume of 10 μL. Amplicons were detected on a Bio-Rad CFX96 thermocycler with the following cycling conditions: 95°C for 3 minutes, followed by 39 cycles of 95°C for 15 seconds and 62°C for 30 seconds. Cycle threshold (Cq) values were determined using the auto-threshold function of the CFX96 software. Each condition was assessed with three biological replicates, with no technical replicates. Parental cells lacking the transgene were included as a negative control; a no-template control using primers alone was also included for the baseline.

### Cell viability assay during differentiation

The parental mESC-Gata4 line and a derivative engineered single clone were used to assess cell viability during differentiation. On Day 0, cells were maintained in 2i medium with or without Shield-1 (16 hours) prior to passaging. On Day 1, cells were seeded at 25,000 cells per well in 96-well plates coated with human laminin and placed under either differentiation medium or 2i medium without Shield-1. Cell viability was assessed at 0, 12, and 24 hours after the Day 1 transfer. Three biological replicates were performed.

At each time point, cells were dissociated into a single cell suspension by aspirating the culture medium, washing once with 1×PBS, and incubating with Accutase at 37°C for 5 minutes. The enzymatic reaction was quenched by adding an equal volume of mESC culture medium, and cells were pelleted by centrifugation at 300×g for 5 minutes. The supernatant was aspirated, and the pellet was resuspended in staining buffer (PBS with 0.1% BSA) at 1 × 10⁶ cells/mL.

To determine cell viability, a working solution of 0.1 μM Calcein AM (Thermo Fisher C3099; 1 mM stock in anhydrous DMSO) was prepared in staining buffer. Cells were incubated for 20 minutes at 37°C in the dark, washed once with 2 mL staining buffer, and centrifuged to remove extracellular dye. The final pellet was resuspended in a 100 μL staining buffer and filtered through a 40 µm filter mesh before acquisition.

Flow cytometry was performed on a Beckman Coulter CytoFLEX S. For data analysis, events were first gated on a forward scatter (FSC) versus side scatter (SSC) plot to exclude debris, followed by an FSC-A versus FSC-H gate to exclude doublets. Viable cells were identified within the singlet gate based on Calcein AM fluorescence (ex/em: 488/525 nm).

### LagTag and CUT&Tag data processing

To demultiplex the resulting sequencing libraries, we generated fastq files from the Aviti sequencer output using the bases2fastq tool (Element Biosciences, version 1.8.0). For the barcoded nb-Tn5 samples, we then assigned reads to each barcode using custom bash scripts making use of basic shell commands and the ‘seqtk’ package (seqtk-v1.5 https://github.com/lh3/seqtk). Briefly, we generated subsetted R1 and R2 fastq files containing reads for each individual designed barcode sequence. We then identified paired R1/R2 reads for each possible combination of 5’ and 3’ barcode sequences and stored these as paired fastq files representing each barcode combination. These paired files were used for downstream data processing.

LagTag and CUT&Tag sequencing data produced in this study were then processed using the nf-core/cutandrun pipeline^71^ (version 3.2.2) with Nextflow^72^ (version 23.10.1). Briefly, sequencing reads were aligned to the mm10 or hg38 reference genome using bowtie2^73^ (version 2.4.4), keeping only reads with a minimum alignment q score of 20. For the barcoded samples, we also removed 42 bp from the 5’ end with parameters --clip_r1 42 --clip_r2 42. BigWig files were generated with defined bin sizes using the bamCoverage function of deepTools^74^ (version 3.5.1) with Counts Per Million mapped reads (CPM) normalization. Similarly, the BigWig files for log2 ratio of LagTag profiles between Lamin B1 and mCherry were generated with defined bin sizes using the bamCompare function of deepTools. The CUT&RUN suspect list regions^75^ were masked. The processed files were then visualized with Integrative Genomics Viewer (IGV)^76^ (version 2.16.2).

### Publicly available datasets

For the mESC analyses, we downloaded ChIP-seq datasets from the ENCODE portal^20,77^ for H3K4me3 (ENCSR212KGS) and H3K27me3 (ENCSR059MBO) and from the NCBI GEO (accession GSE195840)^37^ for RNAPIISer2p. We also used previously called peaks from H3K4me3 ChIP-seq (ENCSR212KGS)^20,77^, H3K27me3 ChIP-seq (https://zenodo.org/records/5507375)^23,56^, and RNAPIISer2p CUT&Tag (accession GSE171554)^23^ by merging peaks from replicates using bedtools (version 2.31.1) intersect^78^. For the Lamin B1 analyses, we downloaded pA-DamID datasets (accession GSE181693)^33^ and lamina-associated domains (LADs) from Peric-Hupkes et al.^57^ for mESCs as well as pA-DamID (4DNESWB729QB)^61^, DamID (4DNES24XA7U8), and TSA-seq (4DNESTE8NJOE)^62^ datasets from the 4D Nucleome data portal^59,60^ for HCT116 cells. The mm9 genomic coordinates were converted to mm10 with the UCSC Genome Browser^79^ program LiftOver. The bigwig files for replicates were averaged with deepTools^74^ (version 3.5.1) bigwigAverage.

### Correlation analysis

Pearson or Spearman correlation coefficients between LagTag (CPM or mCherry log2 ratio normalized), CUT&Tag (CPM), ChIP-seq (input normalized), pA-DamID (Dam-only normalized), DamID (Dam-only normalized), and TSA-seq (input normalized) datasets were calculated with 10-kb binning for chr1-19, X (mESCs) or 20-kb binning (HCT116 cells) using the pearsonr or spearmanr function of scipy.stats^80^ (version 1.11.3). When specified, a subset of 10-kb bins (n = 61,536 out of 251,299 total bins) overlapping the union of peaks previously identified from the ChIP-seq (H3K4me3 and H3K27me3) and CUT&Tag (RNAPIISer2p) datasets in mESCs was used, similar to previous benchmarking^34^. We note that in DamID assays, untethered methyltransferase preferentially labels accessible chromatin^81^, and log2 ratio normalization over a Dam-only control is typically used to correct this bias^14^. We compared CPM and log2 ratio normalized LagTag profiles with orthogonal datasets. Log2 ratio normalization did not improve correlation with orthogonal profiles except for Lamin B1 pA-DamID, suggesting that open chromatin bias does not substantially confound LagTag enrichment signals at the 10-kb resolution used in this study. However, chromatin accessibility bias may pose a greater concern for low-abundance targets such as transcription factors, which were not tested here.

### Signal enrichment

To generate signal enrichment plots, the functions computeMatrix and plotHeatmap from deepTools^74^ (version 3.5.4) were used with the flag --missingDataAsZero. computeMatrix was used in reference-point mode for plots using peak centers and scale-regions mode for plots over gene bodies or LADs. For plots over peak centers or genes, the relevant bigwigs averaged across replicates with a bin size of 50 bp were used, while for plotting for LADs, a bin size of 10 kb was used.

Protein-coding genes were taken from mm10 vM21 annotation (ENCODE accession number: ENCSR884DHJ; filename: mm10_gencode.vM21.primary_assembly.annotation_UCSC_names.gtf). For the gene body heatmaps, these gene annotations were filtered for those with gene_type “protein_coding”.

### Differential enrichment analysis

To perform differential enrichment analysis of RNAPIISer2p signals over gene bodies, deepTools multiBamSummary was first used to generate counts per gene using bam files generated by the cutandrun pipeline^71^. Genes were defined by mm10 vM21 annotation as above but not filtered for protein-coding status. Counts per gene were normalized by dividing by the number of mapped and paired reads per sample. This information was extracted from the cutandrun pipeline results.

Then DESeq2^82^ (version 1.34.0) was used to determine differentially enriched genes with “condition” being the only factor in the design formula. Raw (unnormalized) counts per gene were used as input. The shrinkage method apeglm^83^ (version v1.32.0) was used to shrink log2 fold change estimates. After shrinkage, a gene was deemed differentially enriched between two conditions if the absolute value of log2 fold change exceeded 2 and adjusted P-value was below 0.001.

To generate principal component analysis coordinates, raw counts per gene were first standardized using the DESeq2 variance standardized transformation^82^. Then the prcomp function in R (scale. = FALSE) was run on the transformed counts using the top 500 most variable genes.

The genes displayed in the clustermap were determined by generating a consensus list of differentially enriched genes between conditions 5 and 6 in both the engineered and parental mESC-Gata4 cell lines. The normalized count values over these genes were then processed as follows: the counts were averaged across replicates, then normalized by row (gene) so that each row has a mean of 0 and standard deviation of 1 using the zscore function from scipy.stats^80^. The clustermap function of seaborn^84^ (version 0.11.2) was then used with method “average” to determine row linkage.

### Cell division dilution model

The number of cell divisions after induction of the Shield-1-inducible M.EcoGII fusion protein was estimated using the ratio of sequencing read counts as a linear proxy for variable target abundance (i.e., m6A) relative to the constitutive target (i.e., RNAPIISer2p). Specifically, in LagTag measuring both m6A and RNAPIISer2p targets, the ratio is defined as:

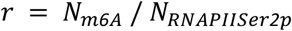

where *N_m6A_* represents read counts from the inducible m6A target, assumed to be diluted by cell division, and *N_RNAPIISer2p_* represents those from the constitutive target. Because RNAPIISer2p target abundance is assumed constant, the ratio *r* is proportional to m6A target abundance. Similarly, the read fraction of m6A antibody was calculated from the same read counts as:

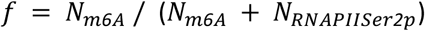

which is related to the ratio by:

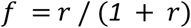

Assuming symmetric partitioning of m6A-marked DNA to daughter cells upon replication^16^, the m6A signal above basal is halved each cell cycle, giving after *n* cell divisions:

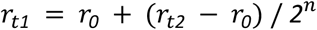

Solving for *n*:

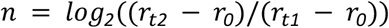

where *r_0_*is the ratio measured in uninduced cells, reflecting background m6A signal in the absence of Shield-1 induction; *r_t2_*is the ratio in cells induced for duration *Δt* ending at timepoint *t2* without no subsequent chase (fully induced reference); and *r_t1_* is the ratio in cells induced for *Δt* before the *t1* timepoint, then chased for (*t_2_* − *t_1_*) without Shield-1 induction. Using this model, we estimated the number of cell divisions for experimental samples at the 24-hour chase timepoint, and computed the predicted dilution curve asymptotically approaching the basal level. To quantify the relative m6A signal abundance above background, enrichment, *E*, was expressed as a normalized fold ratio over the basal level, calculated as:

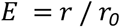

where *r_0_* is the ratio measured in uninduced cells.

### Statistics and reproducibility

CUT&Tag and LagTag experiments were performed with two or three replicates using 96-well plates in Figures 1, 2, and Figures S1-10. No statistical methods were used to predetermine the sample size. The experiments were not randomized and the investigators were not blinded to allocation during the experiments and outcome assessment.

## Supplementary Tables

**Table S1:** A list of primary antibodies used in this study.

**Table S2:** A list of oligonucleotides used in this study.

**Table S3:** Summary of temporal chromatin profiling technologies.

